# Illuminating oncogenic KRAS signaling by multi-dimensional chemical proteomics

**DOI:** 10.1101/2025.01.30.635627

**Authors:** Nicole Kabella, Florian P. Bayer, Konstantinos Stamatiou, Miriam Abele, Amirhossein Sakhteman, Yun-Chien Chang, Vinona Wagner, Antje Gabriel, Johannes Krumm, Maria Reinecke, Melanie Holzner, Michael Aigner, Matthew The, Hannes Hahne, Florian Bassermann, Christina Ludwig, Paola Vagnarelli, Bernhard Kuster

**Affiliations:** School of Life Sciences, Technical University of Munich, Freising, Germany; College of Health, Medicine and Life Science, Brunel University London, London, UK; Bavarian Center for Biomolecular Mass Spectrometry (BayBioMS), School of Life Sciences, Technical University of Munich, Freising, Germany; Department of Medicine III, Klinikum Rechts der Isar, Technical University of Munich, Munich, Germany; TranslaTUM, Center for Translational Cancer Research, Technical University of Munich, Munich, Germany; Deutsches Konsortium für Translationale Krebsforschung (DKTK), Heidelberg, Germany; Bavarian Cancer Research Center (BZKF), Erlangen, Germany; OmicScouts GmbH, Freising, Germany; Partner Site Munich, German Cancer Consortium (DKTK), Munich, Germany

## Abstract

Mutated KRAS is among the most frequent activating genetic alterations in cancer and drug discovery efforts have led to inhibitors that block its activity. To better understand oncogenic KRAS signaling and the cytostatic effects of drugs, we performed comprehensive dose-dependent proteome-wide target deconvolution, pathway engagement and protein expression characterization of KRAS, MEK, ERK, SHP2 and SOS1 inhibitors in pancreatic (KRAS G12C, G12D) and lung cancer (KRAS G12C) cells. Analysis of the resulting 687,954 dose-response curves available online revealed both common and cell line-specific signaling networks dominated by oncogenic KRAS activity. Time-dose experiments separated early KRAS-MEK-ERK from CDK-mediated signaling that cause cells to exit from the cell cycle. This transition to a quiescent state occurred without substantial proteome re-modelling but extensive changes of protein phosphorylation and ubiquitylation. The collective data highlights the complexity of KRAS signaling in cancer and places a large number of new proteins into this functional context.

## Introduction

It is well known that drugs often have more than one target and that the wiring of signaling pathways in cancer cells can be highly diverse leading to sometimes unexpected cellular drug effects (*1–4*). Proteomics approaches have drastically improved our understanding of the molecular and cellular mechanisms of action (MoA) particularly of cancer drugs and selective compounds are increasingly used as chemical probes to study the oncogenic signaling networks these drugs perturb (*5–7*). Understanding the consequences of drug perturbation of the RAS-MEK-ERK axis as one of the most frequently activated pathways in cancer is particularly important. Several proteomics studies using drugs targeting key nodes within the MAPK pathway, such as ERK, MEK, and KRAS, and across different cellular contexts, have reported highly dynamic responses (*8–10*). A notable recent example is a study of the ERK regulated phosphoproteome in KRAS mutated pancreatic cancer cells suggesting that >4,600 phosphorylation sites on >2,100 proteins are directly dependent on ERK activity, implying a broader role of ERK in cancer than hitherto appreciated (*11*).

Given the many functions of ERK for healthy physiology, it is currently unclear if target-related toxicity of ERK inhibition can be adequately managed. In fact, ERK inhibitors have not yet moved beyond phase 1/2 clinical evaluation, in contrast to the approval of currently four MEK and two KRAS inhibitors. The latter have received much attention because oncogenic mutations in KRAS that decouple KRAS activity from upstream signals are detectable in ∼10-20% of all cancer patients (according to The Cancer Genome Atlas database) with G12C being most prevalent in lung and G12D in pancreatic cancer (*12*). The approval of the two KRAS G12C drugs Sotorasib and Adagrasib, which covalently bind Cys12 and trap KRAS in its inactive state, has marked a milestone in KRAS drug discovery (*13*, *14*). Such mutation-specific drugs are attractive because they limit the risk of side effects resulting from target engagement in healthy tissue that also rely on the KRAS pathway for normal function. A number of further modalities are being investigated including inhibitors of KRAS G12D, pan-KRAS inhibitors, compounds directed against active KRAS-GTP, KRAS degraders or drug combinations addressing upstream (EGFR, SHP2, SOS1) or downstream (MEK, ERK) members of the KRAS signaling network that may offer treatment options for a broader range of patients (*15–18*).

However, the cellular MoAs of mutation-specific KRAS inhibitors have not yet been comprehensively characterized on a proteome-wide scale. In the present work, we addressed this gap by measuring the dose-response characteristics of target binding, pathway engagement and proteostasis of Sotorasib, Adagrasib, ARS-1620, MRTX1257 (KRAS G12C inhibitors) and MRTX1133 (KRAS G12D inhibitor), complemented by inhibitors targeting upstream (SHP2, SOS1) and downstream (MEK, ERK) proteins in two KRAS mutated pancreatic (KRAS G12C, G12D) and one lung cancer (KRAS G12C) cell lines. Analysis of the resulting 687,954 dose-response curves highlighted a common core KRAS signaling signature as well as extensive differences between cell lines, placing hundreds of new proteins and their post-translational modifications (PTMs) into the functional context of KRAS signaling. Our data demonstrates that oncogenic KRAS activity dominates the output of MEK and ERK activity, largely decoupled from upstream receptor tyrosine kinase signaling, leading to exit of cells from the cell cycle and transitioning into a quiescent state. Remarkably, it appears that dynamic protein phosphorylation and ubiquitylation rather than protein expression changes are the main drivers of these processes which include inhibition of kinases, transcription factors and E1/E2 ubiquitin ligase activity. We anticipate that the molecular resources provided alongside this manuscript and available in ProteomicsDB.org (*19*) will be of substantial utility for the scientific community for research on the KRAS signaling system and drugs to treat KRAS mutant cancer.

## Results

### Dose-dependent, proteome-wide characterization of KRAS signaling inhibitors

The experimental approach and data compendium created in this study for the analysis of the cellular MoA of mutation-specific KRAS inhibitors and further drugs acting on the RAS-MEK-ERK axis is shown in Figure 1. The distinguishing feature is the systematic measurement of the dose-response characteristics of drug action (characterized by potency (effective concentration to achieve 50% response; EC_50_) and response (curve fold change)) on a proteome-wide scale. Including the EC_50_ dimension is powerful because it enables deducing both, common as well as distinct responses in the same cellular system. We have previously termed this approach decryptE for protein expression and decryptM for PTMs (*5*, *20*). Here, we applied both methods to characterize inhibitors targeting the RAS-MEK-ERK axis and extended the dose-response idea to reactive cysteine profiling for drug-target deconvolution of covalent KRAS inhibitors (decryptC) (fig. S1) (*21*). Statistical analysis of all dose-response data was performed by CurveCurator (*22*) followed by removing outliers or manual inspection if required (S2A; see materials and methods for details). We initially focused on the FDA-approved KRAS G12C inhibitors Sotorasib and Adagrasib, their pre-clinical derivatives ARS-1620 and MRTX1257 and the phase 1/2 KRAS G12D inhibitor MRTX1133 and then expanded to compounds targeting proteins up- and down-stream of KRAS (table S1). For cell line models, we chose the pancreatic cancer lines MiaPaCa-2 (KRAS G12C, homozygous) and ASPC1 (KRAS G12D, homozygous) as well as the lung cancer line NCI-H23 (KRAS G12C, heterozygous). These lines were phenotypically sensitive to KRAS inhibition and represent cancer entities with high clinical prevalence of KRAS G12C or G12D mutations (fig. S3; table S2). Collectively, the different decrypt data types covered 25,038 cysteine-containing peptides (cys-peptides), 69,729 phospho-peptides, 13,093 ubiquitinylated peptides (ubi-peptides) and 8,505 proteins (fig. S1B; table S3-6). Reproducibility of decryptM experiments was assessed by triplicate analysis of phospho-proteomes in response to Sotorasib in MiaPaCa-2 cells showing that 80% of all EC_50_ values and curve fold changes were reproducible within a factor of two (fig. S4) (*5*). All data can be explored in ProteomicsDB (*19*) or via interactive html dashboards provided on Zenodo.org.

**Fig. 1:**
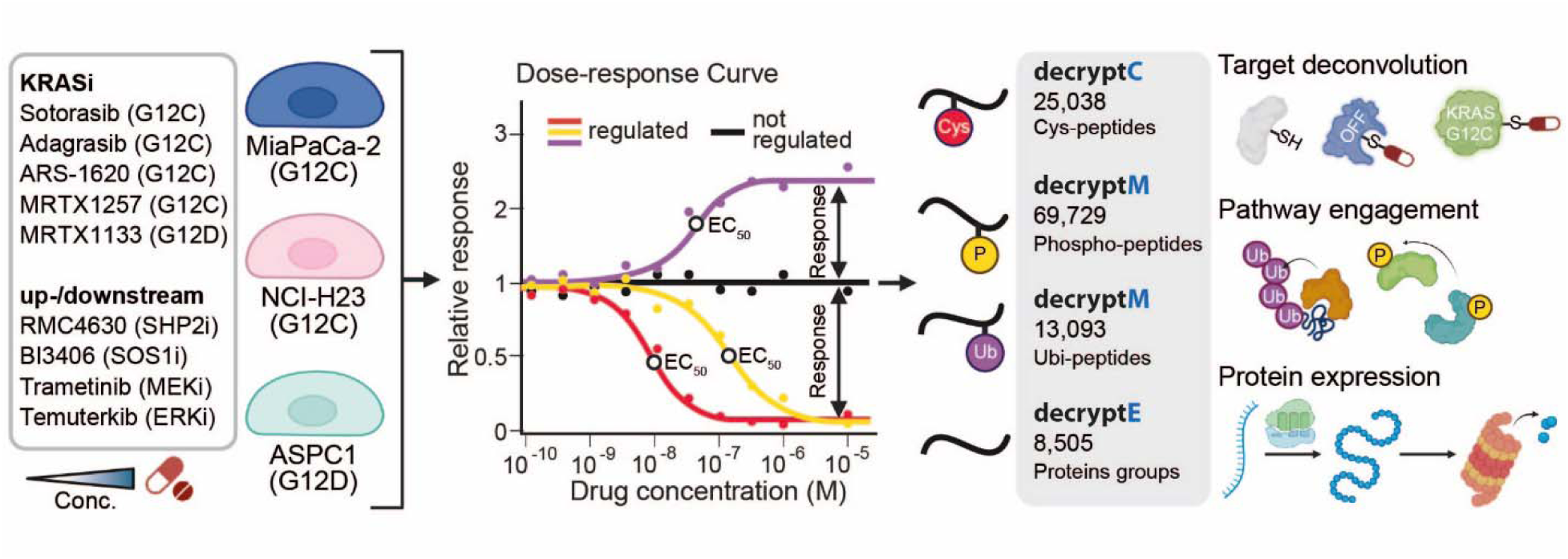
Proteome-wide characterization of KRAS signaling inhibitors by dose-response proteomics. Cells expressing different mutated KRAS proteins were treated with increasing concentrations of the respective drugs and the dose-response profiles of reactive cysteines (for target deconvolution, decryptC), phospho-peptides and ubi-peptides (for pathway engagement, decryptM) and proteins (for protein expression, decryptE) were determined. Drug response (relative to control; up-, down-or not-regulated) and potency (effective concentration to achieve 50% response, EC_50_) were derived from fitted dose-response curves followed by statistical assessment using CurveCurator (see materials and methods for details).

### DecryptC profiling demonstrates high target selectivity of clinical KRAS G12C inhibitors

For *in cellulo* target deconvolution of the KRAS G12C inhibitors Sotorasib and Adagrasib, we applied competitive reactive cysteine profiling in two KRAS G12C and one KRAS G12D (as control) cell lines following 2 hours of drug incubation as previously described (*21*) but extending it here to full dose-response measurements (fig. S1A; table S1; table S3). Between 12,500 and 18,600 cys-peptides were covered per cell line and both G12C inhibitors potently modified C12 of KRAS in the two G12C cell lines (30 nM and 39 nM for Sotorasib, 5nM and 14 nM for Adagrasib in MiaPaCa-2 and NCI-H23 cells, respectively; Fig. 2A,B). These potencies were in line with cell viability data (EC_50_ of 2-7 nM) collected after 72 hours of drug incubation, confirming that the inhibition of KRAS is responsible for the observed phenotypic effect. (fig. S3; table S2). Elongation factor EEF1A2 (C31) was identified as a new but weak (1-3 µM) off-target of Adagrasib, but not Sotorasib in all cell lines (Fig. 2C). Molecular docking provided a rationale for an interaction between EEF1A2-Y86 and the nitrogen of the methylpyrrolidine ring of Adagrasib that does not exist in Sotorasib (Fig. 2D), that stabilizes complex formation between Adagrasib and EEF1A2. Further down-regulated cys-peptides were observed in G12C cell lines which were not regulated in the G12D line. These are likely not *bona-fide* off-targets but result from indirect effects such as loss of protein expression exemplified by two distinct cys-peptides of the transcription factor ETV5 exhibiting equipotent downregulation and similar to KRAS G12C (fig. S5). The collective data shows that Adagrasib and Sotorasib are highly selective for binding to KRAS G12C in cells and thus qualify as chemical probes to study KRAS signaling.

**Fig. 2.**
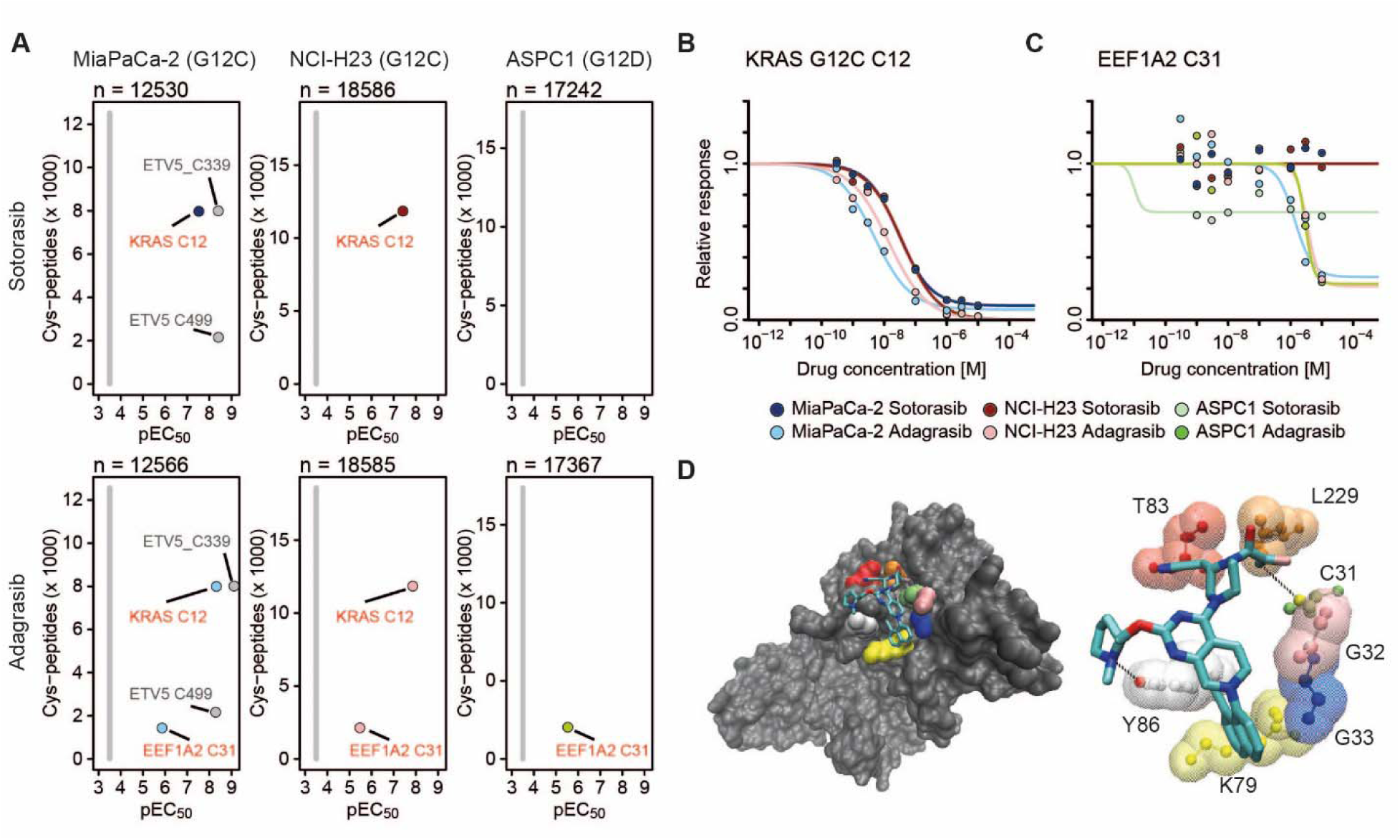
DecryptC profiling of clinical KRASG12C inhibitors in KRAS mutant lines. **(A)** Number of dose-response curves (y-axis) and potency (pEC_50_ = −logEC_50_, x-axis) of cys-peptides following 2 hour treatment of MiaPaCa-2, NCI-H23 and ASPC1 cells with Sotorasib and Adagrasib. Targets of Sotorasib and/or Adagrasib are highlighted in color, all other cys-peptides are displayed in grey. **(B)** Dose-response curves for the primary target KRAS G12C (C12) of Sotorasib and Adagrasib (2 hours). **(C)** Same as (B) but for the Adagrasib off-target EEF1A2. **(D)** Molecular docking of Adagrasib into the structure of EEF1A2. The left panel shows the orientation of Adagrasib within the overall protein structure. The right panel depicts molecular interactions of Adagrasib with amino acid residues of KRAS bringing the reactive enolate group into close proximity to C31 to enable its covalent modification.

### DecryptM profiling demonstrates highly selective pathway engagement of KRAS inhibitors

To examine how mutant KRAS inhibition affects downstream signaling, we subjected Sotorasib, Adagrasib, ARS-1620 and MRTX1257 (targeting KRAS G12C), and MRTX1133 (targeting KRAS G12D) to decryptM profiling of the phosphoproteome using the same cell lines and timing as above (fig. S1A; table S1; table S4). The short treatment time ensured very few changes in protein expression levels (fig. S5A). In KRAS G12C cell lines, KRAS G12C drugs regulated fewer than 600 phospho-peptides (of >20,000 recorded in each experiment; Fig. 3A). In stark contrast, MRTX1133 regulated nearly 2,000 phospho-peptides in ASPC1 (KRAS G12D). Essentially no changes in the phospho-proteome were induced in Sotorasib- and Adagrasib-treated ASPC1 cells, demonstrating high selectivity of KRAS G12C inhibitors for engaging phosphorylation-regulated cellular processes within the chosen drug concentration range. This also means that, the aforementioned Adagrasib off-target EEF1A2 C31 (Fig. 2C,D) had no measurable impact on the phospho-proteome of these cell lines.

**Fig. 3:**
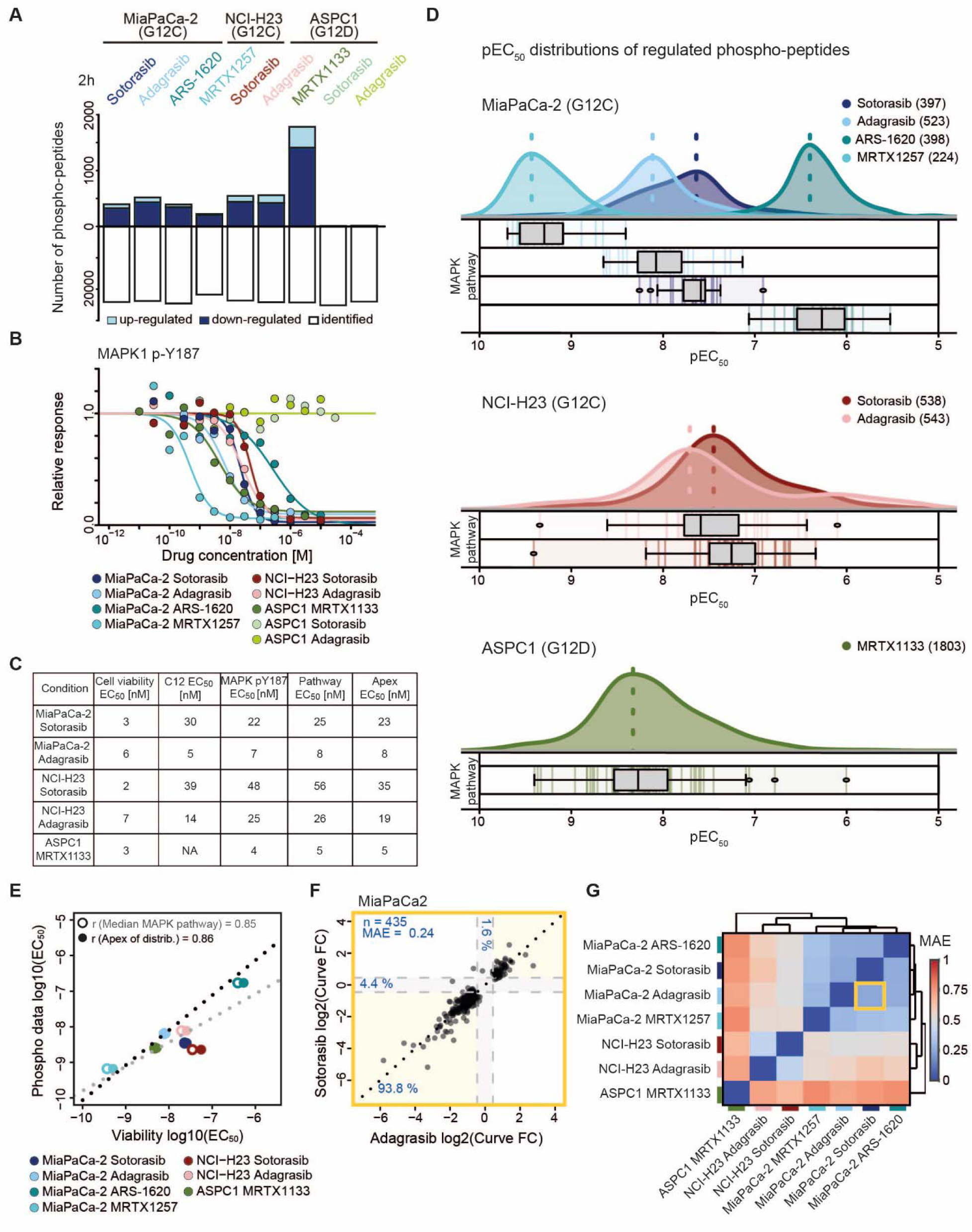
DecryptM profiling of KRAS inhibitors in mutant KRAS cell lines. **(A)** Number of identified and drug-regulated phospho-peptides in three mutant KRAS cell lines after short-term (2 hours) drug treatment. **(B)** Dose-response curves for MAPK1 pY187 activation loop phosphorylation in the same cell lines and using the same drugs as in panel (A). **(C)** Summary of the potency of drug response at the level of cell viability (72 hours), KRAS C12 target binding (2 hours), MAPK1 (Y187) activation loop phosphorylation (2 hours), phosphorylation of KEGG-annotated MAPK pathway members (map04010; median pEC_50_) and all drug-regulated phospho-peptides (apex pEC_50_, 2 hours). **(D)** Upper panels: distribution of drug potency (pEC_50_) determined for all regulated phospho-peptides in a given cell line and treated by a given drug. Numbers behind drug names indicate the number of regulated phospho-peptides detected in the experiment. Lower panels: Boxplots of pEC_50_ values for phospho-peptides on proteins annotated as MAPK pathway members; median indicated by black bar, the box indicates the interquartile range (IQR), its whiskers 1.5× IQR values. **(E)** Correlation analysis of log10(EC_50_) values determined for drug-induced cell viability (72 hours) and phospho-peptide abundance changes (2 hours) either using regulated phospho-peptides from annotated MAPK pathway members only or the apex of the pEC_50_ distribution from all regulated phospho-peptides of each individual treatment (r: Pearson correlation coefficient). **(F)** Scatter plot comparing the log2 curve fold changes of phospho-peptides to Adagrasib and Sotorasib in MiaPaCa-2 cells. Dotted lines mark the boundaries of the applied CurveCurator log2 fold change cut-off (fc-values = ±0.45). Yellow areas contain phospho-peptides with consistent responses, grey areas contain phospho-peptides regulated by one of the conditions only. Percentages indicate the fraction of phospho-peptides in the regions highlighted by color. **(G)** Cluster map summarizing the data exemplified in panel F (orange box) for all KRAS drugs and cell lines based on hierarchical clustering of the mean absolute error (MAE) of log2 curve fold changes.

As expected, KRAS inhibition led to abrogation of MAPK1/3 (ERK) activity, using its activation loop phosphorylation status as a proxy (Fig. 3B). No such effect was observed for KRAS G12C inhibitors in ASPC1 (KRAS G12D). The potency of MAPK1/3 inhibition closely mirrored the decryptC target engagement and cell viability data demonstrating that the drugs fully engaged the KRAS-MEK-ERK axis in cells and are responsible for the observed inhibition of cell growth (Fig. 3C; table S2). The pEC_50_ distributions shown in Figure 3D summarize the dose-response curves of all regulated phospho-peptides for each drug and cell line. It is apparent that drugs had different potencies, with MRTX1257 being the most potent (apex EC_50_ of 0.37 nM) and ARS-1620 the least potent (apex EC_50_ of 398 nM) KRAS inhibitor. Again, these pEC_50_ values closely corresponded to the potency of reactive cysteine profiling (for G12C inhibitors, Fig. 3C) and cell viability data (Fig. 3E). Importantly, within each treatment, many phospho-peptides of known MAPK pathway members (KEGG: map04010; table S7) were regulated within a similar EC_50_ range (Fig. 3D, boxplots). Following a guilt-by-association logic, we conclude that many phospho-peptides with EC_50_ values close to known KRAS pathway members may indeed themselves be functionally linked to the KRAS pathway. To define this group of phospho-peptides, the data was filtered to include only those phospho-peptides that were within ±1 pEC_50_ of the apex of each individual treatment in Figure 3D (fig. S6; see materials and methods for details).

We next asked to what extent phospho-peptides were regulated by KRAS inhibitors in the same or different cell lines. Strikingly, nearly 94% of all phospho-peptides showed consistent regulation in response to Adagrasib and Sotorasib in MiaPaCa-2 cells (Fig. 3F). Pairwise comparison of all drug responses in all cell lines using the mean absolute error (MAE) of the log2 curve fold change as a similarity metric, grouped the data by cell line, not by drug (Fig. 3G; fig. S7). This suggests that the different KRAS drugs act by essentially the same MoA in a given cell line but that the molecular composition and wiring of the KRAS pathway is substantially different between cell lines.

### Identification of a common KRAS core signaling signature connecting multiple cellular processes

The very high target and pathway engagement specificity and selectivity of KRAS inhibitors observed above suggests that the 2,354 short-term (2-hour) drug-regulated phospho-peptides detected in the three cell lines (Fig. 4A) may be members of a wider oncogenic KRAS signaling network. Despite substantial response diversity between the three cell lines, a set of 241 phospho-peptides was regulated in all cell lines regardless of cancer entity, G12C/G12D mutation or hetero-/homozygosity of the KRAS locus (Fig. 4A; fig. S8A; table S8; see materials and methods for details). In the following, we refer to these as the KRAS core signaling signature. These 241 phospho-peptides correspond to 252 phospho-sites on 196 proteins (canonical sequences of protein coding genes, fig. S8B). We compared our data at the phospho-site level to the ERK compendium (*8*), a previously published study focused on the ERK-dependent phospho-proteome (*11*), and all regulatory phospho-sites documented in PhosphoSitePlus (phosphosite.org). Despite a very large overlap of our data to these resources in terms of phospho-proteome coverage (71% overlap with the ERK compendium, 87% with the ERK-dependent phospho-proteome and 22% with PhosphoSitePlus; fig. S8C), the number of shared drug-regulated phospho-sites was relatively low (Fig. 4B). This discrepancy may be attributed to several factors: i) other resources called regulation on the basis of p-values from replicate experiments of high single-concentration drug treatments, whereas we employed dose-response statistics; ii) not all cell lines used were the same; and iii) the drug treatment times were not identical. Still, 67% of the KRAS core signaling signature defined above was also contained in the ERK compendium and the ERK-dependent phosphoproteome, underscoring its robustness (Fig. 4B).

**Fig. 4:**
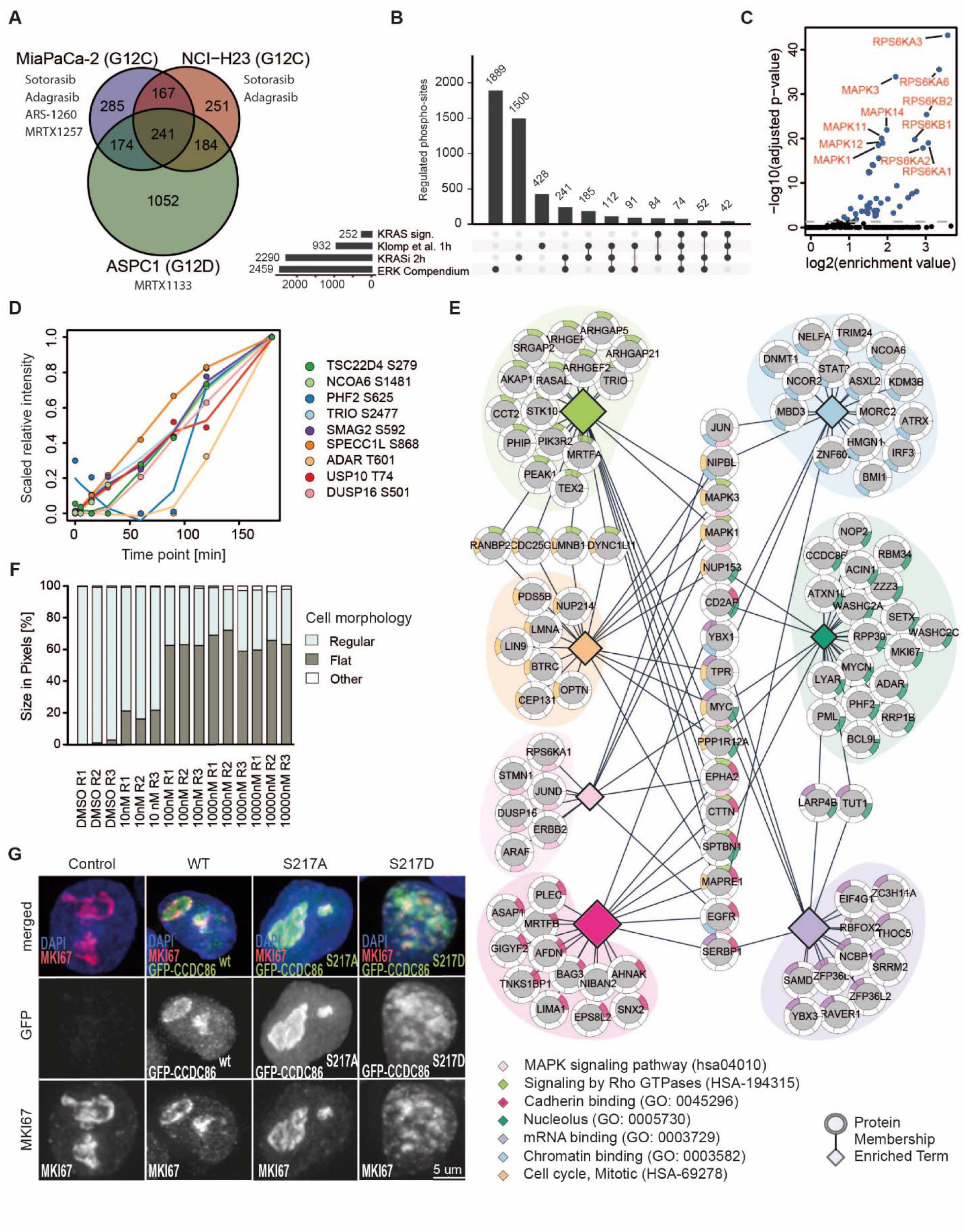
Cellular processes impacted by a KRAS core signaling signature. **(A)** Venn diagram of drug-regulated phospho-peptides detected in MiaPaCa-2 (KRAS G12C), NCI-H23 (KRAS G12C) and ASPC1 (KRAS G12D) cells defining a common KRAS core signaling signature of 241 phospho-peptides. **(B)** Upset plot comparing data from the present study (252 phospho-sites on the 241 phospho-peptides that comprise the KRAS core signaling signature and all 2,290 phospho-sites on the 2,354 phospho-peptides regulated by all KRAS drugs after 2 hours of treatment) to two published molecular resources of ERK signaling (the ERK-regulated phospho-proteome, denoted as Klomp et al. and the ERK compendium (*8*, *11*)). **(C)** Kinase motif enrichment analysis of down-regulated phospho-peptides from the KRAS core signaling signature. **(D)** Results of MAPK1 *in vitro* kinase assays using synthetic peptides representing putative MAPK1 substrates from the list of 241 phospho-peptides comprising the KRAS core signaling signature. For clarity, the response of the assay was scaled from 0 to 1. **(E)** Graphical representation of enriched biological processes of proteins underlying the KRAS core signaling signature. The size of the diamond node for each enriched term was scaled by the statistical significance of the enrichment analysis (adjusted p-val.). Protein nodes with more than one color in the halo map to more than one term. **(F)** Quantification of cell morphology features of MiaPaCa-2 cells in response to Sotorasib (72 hours). **(G)** Representative images of NCI-H23 cells untransfected (control) or transfected with either GFP–CCDC86 WT, GFP–CCDC86 S217A or GFP–CCDC86 S217D constructs (green) after 24 hours. Cells were also stained for MKI67 (red) and DNA (DAPI). Scale bar 5 µm.

Kinase motif enrichment analysis (*23*, *24*) (phosphosite.org) of the KRAS core signaling signature highlighted ERK (SP/TP motif) and several members of the RSK, MNK and S6K families (basophilic motif) as the major kinases that likely phosphorylate these peptides (Fig. 4C; table S7). This is supported by the observation that several phospho-sites known to regulate the activity of these kinases were inhibited by the KRAS drugs (*25*)(according to phosphosite.org; Fig. 3B; fig. S9A). The very same kinases also dominated when performing kinase motif enrichment analysis with all short-term (2-hour) drug-regulated phospho-peptides (fig. S9B; table S7). About 51 % (n=124) of the peptides in the KRAS signaling signature were phosphorylated at a SP/TP site and several of these are annotated substrates of MAPK1 and MAPK3 including the transcription factor ERF (p-T562; fig. S9C)(*26*). Given the pronounced inhibition of MAPK1/3 activation loop phosphorylation by KRAS inhibitors, we hypothesized that many of the hitherto uncharacterized SP/TP motif-containing phosphorylation sites may be novel MAPK1/3 substrates. To test this, we performed in-vitro kinase assays using recombinant MAPK1, synthetic peptide substrates and monitored phosphorylation by a time-resolved parallel reaction monitoring (PRM) assay (table S9). Nine of the 19 tested candidates showed increased phosphorylation over time (Fig. 4D). Among the underlying proteins were the transcriptional regulator SMAG2 (S592), the nucleotide exchange factor TRIO (S2477) that controls RHO and RAC1 activity and ADAR (T601), a protein involved in RNA editing. Interestingly, USP10 (T74), a ubiquitin hydrolase that removes ubiquitin from target proteins such as p53 and CFTR (*27*, *28*), was also among the potential novel direct MAPK1/3 substrates. In addition, for some of the new MAPK1/3 substrates (SPECC1L S868, TSC22D4 S279, ADAR T601 and PHF2 S625), previous studies have already indicated dependency on the activation state of the MAPK pathway and our kinase assays now provides direct evidence that MAPK1/3 can be the phosphorylating kinase (*10*, *29*, *30*).

STRING protein interaction analysis of the 196 proteins underlying the KRAS core signaling signature revealed that about half of these proteins were not connected to each other or to KRAS (fig. S9D). Yet our data tightly connects them to the KRAS network. Many of these phospho-proteins have diverse or poorly characterized functions suggesting that KRAS signaling extends far beyond well-researched biology and that such new avenues may be discovered on the basis of the data provided here. Functional enrichment analysis of the same proteins highlighted seven categories comprised of 106 proteins (Fig. 4E; table S7). Expectedly, these included well-known members of the MAPK pathway such as MAPK1/3 (p-Y187/p-Y204), DUSP16 (p-S501) and ARAF (p-T181) as well as the receptor tyrosine kinases (RTK) EGFR (p-T693), ERBB2 (p-T701, p-S1054) and EPHA2 (p-S901). The latter represent described feedback signaling that is associated with resistance to KRAS inhibition (*31–33*). Important additional functional links could be established to Rho signaling, chromatin binding, mRNA binding, cadherin binding, mitotic cell cycle and localization to the nucleolus. Hence, the 2-hour drug treatment data already illuminates phenotypic drug responses long before they manifest visually under a microscope. This is exemplified by changes in cell shape in response to Sotorasib or MRTX1133 in MiaPaCa-2 or ASPC1 cells after 72 hours (Fig. 4F; fig. S10A,B; table S2) which aligns with the observed regulation of phospho-sites on several proteins involved in Rho signaling (*34*). Gene set enrichment analysis also uncovered a link to the nucleolus including altered phosphorylation levels of the cell proliferation marker MKI67 (p-S1533) and its recently identified interaction partner CCDC86 (p-S217)(*35*). Immuno-fluorescence microscopy showed that KRAS inhibition (24 hours) led to the loss of MKI67 signal indicating loss of nucleolar integrity in MiaPaCa-2 and NCI-H23 cells (fig. S10C-E; table S2). To investigate if CCDC86 S217 phosphorylation may be functionally important, we expressed GFP-tagged CCDC86 WT, S217A and S217D in NCI-H23 cells. While cells expressing wild type or an Alanine mutant of the protein showed no phenotype, expression of the phospho-mimetic CCDC86 S217D drastically disrupted nucleolar integrity in 25% of cells 24 hours post-transfection (Fig. 4G; fig. S10F,G; table S2). The mechanism by which this happens is currently not clear but this very strong phenotype suggests that the phosphorylation status of CCDC86 S217 is of critical importance for nucleolar integrity and that this phospho-site is responsive to KRAS drugs.

### Mutant KRAS activity dominates the signaling output of MEK and ERK

We next expanded decryptM profiling (2 hours) to inhibitors that act on targets upstream (SOS1, BI3406; SHP2, RMC4630) or downstream (MEK, Trametinib; ERK, Temuterkib) of KRAS (fig S2B; table S1, table S4) to investigate two main questions. First, does the additional data provide independent support for the KRAS core signaling signature? And yes, a total of 57% (NCI-H23), 80% (MiaPaCa-2) and 86% (ASPC1) of the KRAS core signaling signature was also regulated by the additional drugs (fig. S11). Second, to what extent does hyper-active KRAS activity alone determine the output of downstream MEK and ERK activity independent of upstream input signals provided by receptor tyrosine kinase (RTKs) activity? Answering this question requires that the employed chemical probes are highly selective for their targets. This was already established above for the KRAS inhibitors. The decryptM profiles of RMC4630, Temuterkib and Trametinib, however, showed bi-modal pEC_50_ distributions implying off-target/off-pathway effects (fig. S11). The ability of decryptM profiling to recognize such off-target/off-pathway effects, highlights the superiority of the dose-response approach over traditional experiments using a fixed, and often arbitrarily high drug concentration (fig. S12, S13A). Kinase motif enrichment suggested, and Kinobead selectivity profiling confirmed, AAK1 (K_d_ ^app^ 11 nM) and GAK (K_d_ ^app^ 155 nM) as off-targets of Temuterkib (fig. S13A-C; table S7,10) but no off-targets were found for Trametinib (*1*). To ensure high quality of the subsequent analysis, drug-regulated phospho-peptides were filtered such that they follow the pEC_50_ distribution of KRAS core signaling signature members (Fig. 5A; fig. S12; see materials and methods for details).

**Fig. 5:**
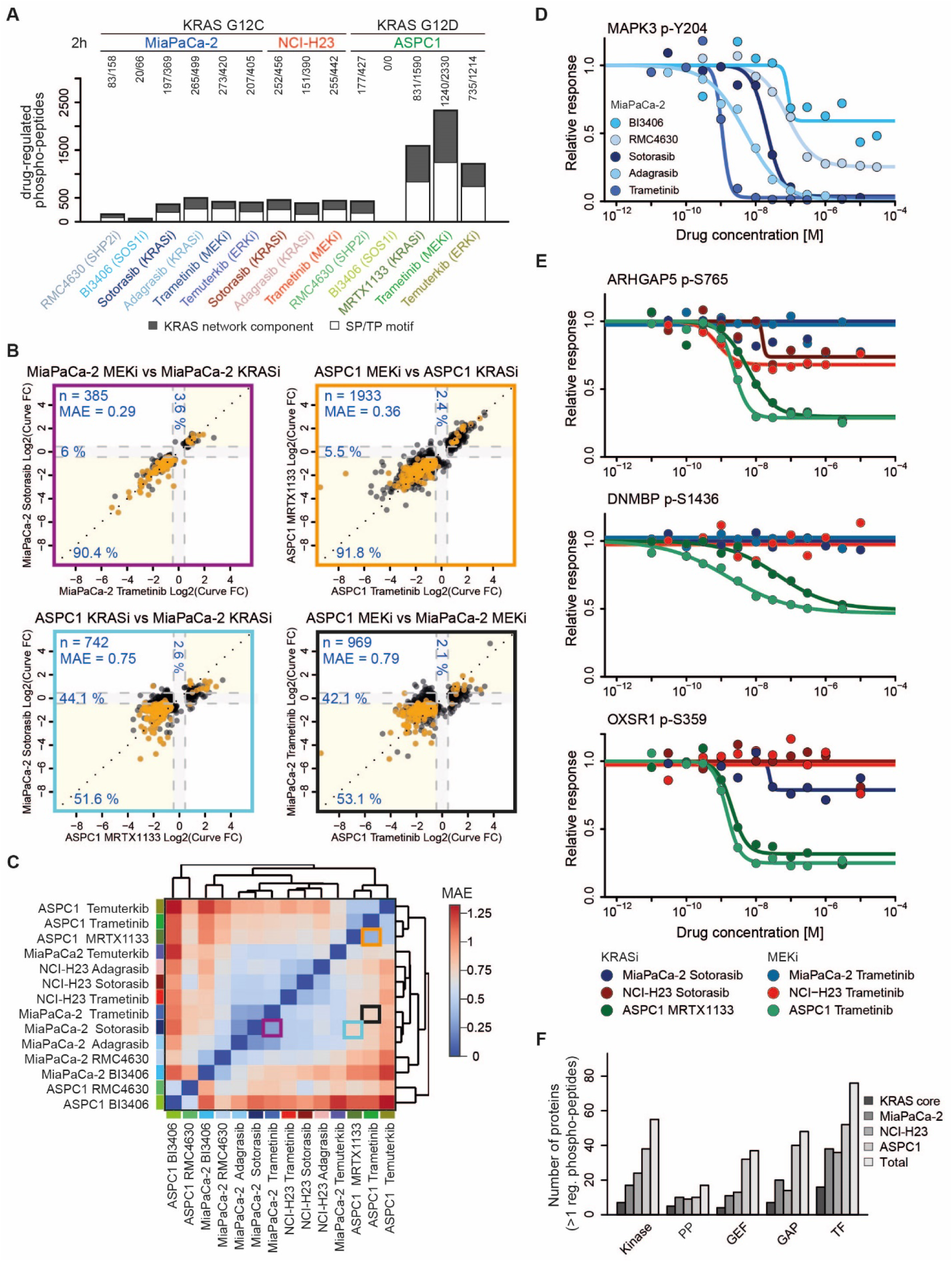
Cell line-specific KRAS signaling. **(A)** Number of drug-regulated phospho-peptides in response to BI3406 (SOS), RMC4630 (SHP2), Sotorasib (KRAS), Adagrasib (KRAS), MRTX1133 (KRAS), Trametinib (MEK) and Temuterkib (ERK) after 2 hours of drug exposure in three mutant KRAS cell lines. The black part of each bar represents the number of phospho-peptides containing an SP or TP phosphorylation motif. Numbers on the top of each bar specify the number of phospho-peptides in the white and black parts of a bar respectively. **(B)** Pairwise comparisons of drug responses (log2 curve fold change) of phospho-peptides for different combinations of drugs and cell lines. Yellow areas contain phospho-peptides with consistent responses, grey areas contain phospho-peptides regulated by one of the conditions only. Dotted lines mark the boundaries of statistical significance based on a CurveCurator log2 fold change cut-off (fc-values = ±0.45). Percentages indicate the fraction of phospho-peptides in the respective colored regions and n denotes the number of phospho-peptides in the plot. Orange data points mark phospho-peptides from the KRAS core signaling signature. MAE: mean absolute error of log2 curve fold changes. **(C)** Cluster map summarizing the analysis shown in (B) for all combinations of drugs and cell lines from (A). **(D)** Dose-response curves of MAPK3 Y204 phosphorylation (activation loop) in response to 7 MAPK pathway modulating drugs. **(E)** Example dose-response curves of phospho-peptides that were significantly drug-regulated in one cell line (here ASPC1) but not in others. **(F)** Number of kinases, phosphatases (PP), guanine nucleotide exchange factors (GEFs), GTPase-activating proteins (GAPs) and transcription factors (TF) that showed drug-induced regulation of at least one phospho-peptide in the different cell lines upon KRAS inhibition.

Comparison of decryptM data collected for MiaPaCa-2 and ASPC1 cells in response to KRAS and MEK inhibition revealed a striking consistency of phospho-proteome responses (90.4% and 91.8% respectively, Fig. 5B). Similarly strong consistencies were observed for KRAS and ERK inhibition (81.8% and 89.9% respectively; fig. S14). Hierarchically clustering the results of these pair-wise comparisons separated SOS1 and SHP2 inhibitor effects from those of KRAS, MEK and ERK inhibitors (Fig. 5C; fig. S14). In addition, unlike KRAS and MEK inhibition, SHP2 and SOS1 inhibition did not fully abrogate downstream MAPK3 (p-Y204) activity in MiaPaCa-2 cells (Fig. 5D; fig. S15). Particularly SOS1 inhibition showed only weak pathway engagement illustrated by the regulation of only few phospho-peptides in MiaPaCa-2 and none in ASPC1 cells (Fig. 5A). Consequently, cellular assays showed that neither SHP2 nor SOS1 inhibition had a substantial effect on cell viability (fig. S3; table S2). Together, this suggests that mutant KRAS largely (MiaPaCa-2) or almost completely (ASPC1) decouples the MAPK pathway from upstream signals and that mutant KRAS activity, therefore, dominates the signaling output of the downstream kinases MEK and ERK.

### Cell line-specific wiring of mutant KRAS signaling

The remarkable consistency of phospho-proteome responses between KRAS vs. MEK and KRAS vs. ERK inhibition not only applied to the KRAS core signaling signature but also included many more phospho-peptides within the same cell line. As a result, the mutant KRAS-controlled signaling network may extend beyond 800 potential members in MiaPaCa-2 cells, 700 in NCI-H23 cells, and 1,900 in ASPC1 cells (fig. S16; table S8). At the same time, comparing decryptM profiles between cell lines showed that the response of the same phospho-peptide to inhibition of the same target protein can be very different. This is evident from comparing the consistency of phospho-peptide regulation induced by the same drug but in different cell lines. For instance, consistency for KRAS or MEK inhibition in ASPC1 vs. MiaPaCa-2 cells (51.6% and 53.1% respectively) was much lower than KRAS vs. MEK inhibition in MiaPaCa-2 and ASPC1 cells (> 90%; Fig. 5B,C; fig. S14). ASPC1 cells exhibited a particularly distinct response to KRAS, MEK, and ERK inhibition and many phospho-peptides showed consistent drug-regulation for all three drugs, while several of these were never even detected in any other cell line (fig. S16B). More interestingly, many phospho-peptides were clearly drug-regulated in ASPC1 cells but not in any other cell line despite the fact that they were detected (Fig. 5E). Functional enrichment analysis of drug-regulated phospho-peptides performed separately for each cell line highlighted biological functions already found for the KRAS core signaling signature. For instance, Rho signaling was independently enriched in the data of all three cell lines but a particularly large number of phospho-peptides on related proteins were identified in ASPC1 cells suggesting a stronger connection between KRAS signaling and the regulation of the actin cytoskeleton in ASPC1 compared to the other cell lines (fig. S16C; table S7). Similarly, the top 15 Kinase motifs were enriched similarly strongly in each cell line even though the underlying phospho-peptides were not necessarily the same (mainly MAPK and RPS6K motifs) even though the underlying phospho-peptides were not necessarily the same (fig. S16D, table S7). Despite substantial differences in the absolute number of regulated phospho-peptides, the proportion of proline-directed motifs (SP/TP; associated with MAPKs) was similar between the cell lines (Fig. 5A). In light of this data, we propose, that although the same underlying molecular mechanisms (i.e. inhibition of the RAS-MEK-ERK axis) drive the observed regulation of phospho-peptides in each cell line, the particular molecular wiring of that network depends on the presence, abundance and activity of cell line-specific factors such as certain kinases, phosphatases, guanine nucleotide exchange factors, GTPase-activating proteins or transcription factors (Fig. 5F).

### Two-dimensional decryptM distinguishes immediate KRAS pathway inhibition from adaptive cellular response

KRAS inhibition was cytostatic in all tested cell lines but did not induce cell death (fig. S3; fig. S10). To shed light on the steps leading to this cellular adaptation, we added a kinetic component to decryptM profiling of Sotorasib in MiaPaCa-2 cells (dose-dependent treatment for 1, 2, 8, and 16 hours; table S1). Between 271 and 401 phospho-peptides showed dose-dependent changes during the first 8 hours increasing to 1,075 after 16 hours (fig. S17A; table S4). We classified the 550 phospho-peptides that were detected at all four time points into four categories (Fig. 6A; table S1; see materials and methods for details): The constant group (n=231) showed drug regulation at three or all four time points but always at 16 hours (e.g. the activity-inducing p-S380 site on RPS6KA1) thus representing drivers of sustained drug response. The early group (n=50) showed regulation at 1 hour and/or 2 hours and/or 8 hours but not at 16 hours, possibly representing initiators of cellular adaptation, exemplified by BRD3 p-S281. For the sake of clarity, we combined these two groups to describe the immediate response of cells that is a direct consequence of KRAS inhibition. The intermediate group (n=69) showed regulation at 8 hours and 16 hours only, exemplified by the E3 ubiquitin ligase RNF168 p-T208 and the late group (n=200) showed regulation at 16 hours only and included the cell cycle regulator WEE1 (p-T190). The two latter categories most likely represent the result of the adaptive response and were, therefore combined. t-SNE analysis nearly completely separated the immediate and adaptive responses (Fig. 6B). Practically all phospho-peptides of the KRAS core signaling signature belonged to the immediate response group (Fig. 6C). In contrast to a recently published study (*11*), this clear temporal distinction of drug-regulated phospho-peptides indicates that not all are the direct consequence of KRAS inhibition.

**Fig. 6:**
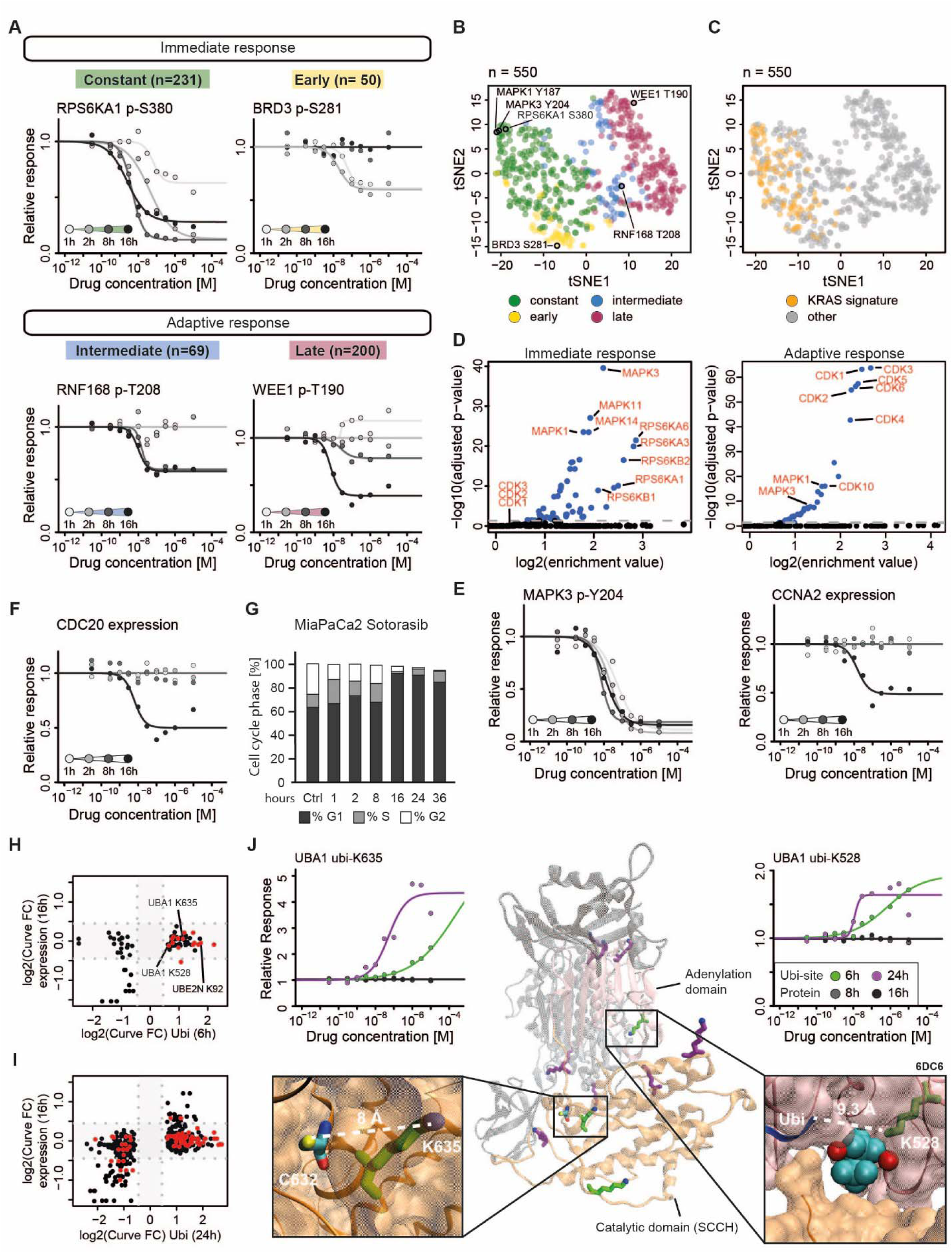
Temporal response of the proteome, phospho-proteome and ubiquitinome of MiaPaCa-2 cells to Sotorasib. **(A)** Example dose-response curves for phospho-peptides that showed constant (green), early (yellow), intermediate (blue) or late (red) response to 1-16 hours of Sotorasib treatment. **(B)** t-SNE plot of responses (abs. log2 curve fold change) of phospho-peptides detected at all four time points and colored by the groups defined in (A). n denotes the number of phospho-peptides in the plot. **(C)** Same as (B) but highlighting members of the KRAS core signaling signature. **(D)** Kinase motif enrichment analysis of down-regulated phospho-peptides from the constant/early groups (immediate response; left panel) or intermediate/late groups (adaptive response; right panel). **(E)** Dose-response curves of MAPK Y204 phosphorylation (indicating kinase activity) and Cyclin A (CCNA2) protein abundance (indicating CDK activity) at the four time points. **(F)** Dose-response curve of protein abundance of the cell cycle regulating protein CDC20. **(G)** Barplot summarizing FACS-based cell cycle analysis data (propidium iodide staining) following different durations of Sotorasib treatment in MiaPaCa-2. **(H+I)** Scatter plot comparing protein abundance changes (16 hours) of proteins that show drug-regulated ubi-peptides at 6 hours and 24 hours respectively. Proteins of the ubiquitin conjugation system are highlighted in red. Dotted lines mark boundaries of statistical significance based on the applied CurveCurator log2 fold change cut-off (fc-values = ±0.45). Grey areas contain ubi-peptides or proteins regulated by one of the conditions only **(J)** Alphafold structure of UBA1 (*46*) (center panel) highlighting the adenylation domain (pink) and catalytic SCCH domain (orange) as well as drug-regulated ubiquitinylated lysine residues after 6 hours (green) or 24 hours (purple) of Sotorasib. Left top panel: dose-response curves of UBA1 K635 ubiquitylation after 6 hours (light green) and 24 hours (purple) of Sotorasib. Left bottom panel: Magnified view of the catalytic domain highlighting drug-regulated ubiquitinylated K635 (green) in close proximity to the catalytic C632 residue based on Alphafold structure of UBA1. Right top panel: dose-response curves of UBA1 K528 ubiquitylation after 6 hours (light green) and 24 hours (purple) of Sotorasib. Right bottom panel: magnified view of the co-crystal structure of UBA1 (PDB: 6DC6) (*47*), ATP(β,γ)+Mg (cyan and red) and ubiquitin (blue) highlighting drug-regulated ubiquitinylated K528 in close proximity.

As one might expect, kinase motif analysis for the immediate response showed enrichment for ERK as well as members of the RSK family (Fig. 6D; table S7). This is reflected by the full inhibition of activation loop phosphorylation of MAPK3 (p-Y204) at all time points (Fig. 6E). In contrast, the adaptive response enriched for motifs phosphorylated by the cell cycle-regulating kinases CDK1-6 (Fig. 6D; table S7). MAPKs and CDKs both phosphorylate SP/TP sites and we observed a shift in the population of drug-regulated SP/TP-containing phospho-peptides starting from 48% at 1 hour, increasing to 55% at 2 hours and 8 hours and reaching 75% at 16 hours of treatment (fig. S17A). In parallel, we observed down-regulation of protein abundance of Cyclin A (CCNA2; Fig. 6E, table S5) and CDC20 (relevant for M-phase progression; Fig. 6F) as well increased protein abundance of the cell cycle inhibitor CDKN1B (p27; relevant for G0/G1) indicating exit from cell cycle (fig. S17B). FACS analysis of drug-treated MiaPaCa-2 (Fig. 6G) and NCI-H23 (fig. S17C) cells confirmed a decrease of cells in S- and G2-phase following 16 hours of drug treatment and a concomitant increase of cells in G1 (table S2). Therefore, we conclude that the changes in phosphorylation levels in the adaptive response reflect the depletion of active CDK1-3 as a result of a shift in the underlying cell population rather than being a direct consequence of RAS-MEK-ERK signaling inactivation. Collectively, this also suggests that the phospho-peptides of the constant group are the functionally most important triggers of the adaptive cellular response which manifests by cells eventually exiting the cell cycle, entering a quiescent state, thus evading cell death.

### Post-translational protein modifications are the primary regulators of cellular adaptation to KRAS inhibition

Cells changing from a proliferative to a resting state need to down-regulate many cellular processes in one concerted action and many of these could be traced in the 2D-decryptM data. Interestingly, only very few statistically significant changes were observed at the protein level (16 after 8 hours; 53 after 16 hours, fig. S17D; table S5) of the ∼7,000 proteins monitored in the experiment even though the majority of cells had already arrested (Fig. 6G, fig. S17C). Down-regulated proteins included ∼10 transcription factors (including MYC, JUN, FOSL1 and SOX9) implying reduction of target gene transcription as described earlier (*36*), but without numerically large changes in protein levels of their target genes (fig S17B). Five cell cycle regulating proteins (CCNA2/Cyclin A2, CDC20, CDCA5 and UHRF1), and several negative regulators of RTK-MAPK signaling axis (SPRY2/4, SPRY2, ERRFI1) were also down-regulated. Even fewer proteins showed a clear dose-dependent upregulation of protein abundance most notably the CDK inhibitor CDKN1B as well as AGO1, a key-player in post-transcriptional gene silencing, again pointing to cell cycle arrest and down-regulation of transcription.

Our data strongly suggests that cellular adaptation to KRAS inhibition is primarily mediated by PTMs rather than protein expression. For example, dozens of phospho-peptides from chromatin-modifying enzymes (Reactome HSA-3247509) and DNA repair proteins (Reactome HSA-73894) were regulated in both, the immediate and adaptive responses (fig. S17E, table S7). The former includes several lysine acetyl transferases, methyltransferases and demethylases - enzymes that control the state of chromatin activity. Similarly, loss of phosphorylation on proteins important for DNA repair (such as BRCA2 p-S93, ATR p-S435 or TP53 p-S315) likely reflects reduced requirements for DNA repair activity in non-dividing cells. Many and often large changes in PTMs were also detected at all the levels that regulate proteostasis. Reduction in transcriptional activity leads to reduction of mRNA processing activity and we observed regulation of several phospho-proteins associated with this process such as NCBP1 (p-T21) and THOC5 (p-T328) (Fig. 4E). Similarly, changes in phosphorylation were detected on proteins important for translation (e.g. EIF4G1 p-S1231; Fig. 4E).

Given the critical role of the ubiquitin system for cellular proteostasis, we investigated the ubiquitylation status of the proteome in response to 6 and 24 hours of KRAS inhibition by Sotorasib in MiaPaCa-2 cells. This analysis revealed ∼800 ubi-peptides (of 13,000 monitored; fig. S18A-C; table S6) that were dynamically regulated by Sotorasib, mostly independent of their protein levels (Fig. 6H,I). We observed a dose-dependent increase in ubiquitylation levels of many proteins involved in ubiquitin conjugation as early as 6 hours, notably UBA1 (e.g. ubi-K802, ubi-K528, ubi-K657), one of only two ubiquitin-activating (E1) enzymes found in humans (Fig. 6J). Interestingly, the earliest changes occurred in both the catalytically active adenylation domain (AAD) as well as the catalytically active SCCH domain responsible for thio-ester bond formation, both critical for ubiquitin activation (*37*). The ubiquitin-modified lysine residues (ubi-K528 and ubi-K635) are located inside the active sites of UBA1, possibly reflecting an auto-ubiquitylation mechanism and leading to attenuation of enzymatic activity. Similarly, increased ubiquitylation was also observed near the catalytic residues of several E2 enzymes as well as residues that mediate interactions with E1 and E3 enzymes (e.g. UBE2N, UBE2S, and UBE2L3; fig. S18D,E). Lysine ubiquitylation near active site cysteine residues of E2 proteins was shown to impact protein function of the E2 protein UBE2S before (*38*) and a recent study suggested that such E1 and E2 ubiquitylation occurs by ‘random’ E3 ligase activity (*39*). However, our data clearly shows that the ubiquitylation events on E1 and E2 enzymes are dynamically regulated upon KRAS inhibition and the resulting cell cycle arrest. This suggests that attenuating the activity of the ubiquitin system is another and controlled step in the concerted action of cells to transition from a proliferative to a quiescent state.

### Conclusions

KRAS has long been considered to be a critical driver of many cancers and the approval of Sotorasib and Adagrasib in 2021 and 2022 respectively have expanded the therapeutic options for patients with KRAS G12C-mutated, locally advanced or metastatic non-small cell lung cancer (NSCLC). Unfortunately, neither mutant KRAS inhibition nor degradation (*15*) typically kill cancer cells and clinical studies combining KRAS drugs with inhibitors targeting up-or downstream proteins (including EGFR, SHP2, SOS1, MEK and ERK) have also not yet achieved tumor cell eradication (NCT04975256, NCT04185883, NCT0418588 (*40*)). This study aimed to shed new light on the effects of KRAS inhibition by performing extensive, multi-dimensional and fully dose-resolved chemical proteomics in three KRAS mutant lung and pancreatic cancer cell lines. We demonstrated that KRAS inhibitors are highly selective which allows using them as chemical probes to study mutant KRAS-dependent signaling. The phospho-proteome data showed that KRAS mutations strongly decouple up-from downstream signaling such that SHP2 or SOS1 inhibition has little to no effect on the KRAS pathway in KRAS mutant cell lines. Because the phospho-proteome responses of ERK and MEK inhibition were largely indistinguishable from KRAS inhibition, we conclude that the fate of these cancer cells is decided at the level of KRAS inhibition. The drugs and cellular models used here also enabled the definition of a KRAS core signaling signature with many new molecular players that operates in all cell lines and impacts the same major cellular processes.

Importantly, we observed a clear distinction between immediate drug responses, predominantly reflecting KRAS signaling, and adaptive drug responses, which result from cells exiting the cell cycle. The anti-proliferative effect of KRAS inhibition is driven by the immediate regulation of PTMs, leading to the down-regulation of proteostasis and inhibition of cell cycle entry, which results in the observed accumulation of quiescent cells. This transition is mediated by changes on PTM-level rather than protein expression, and may not only allow cells to avoid the energy costs associated with substantial protein expression re-modeling, but may also enable cancer cells to return rapidly to a proliferative state when conditions improve. From a drug discovery point of view, this suggests that mutant KRAS inhibition or degradation may have to be combined with drugs that prevent the escape mechanism via exit from the cell cycle in order to kill the cancer cell. A very recent report suggests that targeting WEE1, PLK1 or CHK1 may be promising in this regard (*41*) and the molecular resource created by the current study (available in ProteomicsDB (*19*)) and in the form of interactive dashboards, may help scientists to identify further such vulnerabilities for future exploitation.

## Materials and methods summary

Full details of the methodology are described in the supplementary materials and are summarized here as follows: Cell lines were treated with increasing concentrations of a drug for a fixed amount of time. Cells were then lysed, proteins extracted, and peptides generated by trypsin digestion. The peptides were stable isotope–labeled using TMT (*42*). Post-translationally modified peptides (phospho-peptides and ubi-peptides) and cysteine-containing peptides (cys-peptides) were enriched by affinity-based methods. Full proteome, cys-peptides or PTM samples were analyzed by LC-MS/MS, and peptides and proteins identified and quantified using MaxQuant/Andromeda (*43*, *44*) or ProteomeDiscoverer. Dose-response curves for each peptide were fitted to a sigmoidal curve model using CurveCurator (on the basis of TMT reporter ion intensities)(*22*), yielding drug EC_50_ values for each protein, cys-or PTM peptide, the size of each effect (curve fold change), as well as curve-fitting quality metrics. A set of filters was applied to extract regulated dose-response curves from each experiment. The dose-dependent regulated PTMs, cysteines or protein expression values were further analyzed to explore the MoA of the respective drugs and the specific responses of different cell lines. Regulated curves were functionally annotated using several knowledge bases.

## Acknowledgments

We thank Kara Kreutz for contributions to sample processing and Karl Kramer for analysis of phenotypic data. Parts of figures were created using BioRender.com.

## Author contributions

NK, and BK conceived the study. N.K., K.S., YC.C., M.A., V.W., A.G., M.R., M.Ai., M.H., F.P.B. performed laboratory experiments. N.K., A.S., K.S., M.A., M.R. and B.K. performed data analysis. J.K., P.V., C.L., H.H., M.T., F.B. and B.K. directed and supervised experiments and data analysis. N.K. and B.K. wrote the manuscript with input from all authors.

## Data and materials availability

The data supporting the findings of this study including raw mass spectrometry proteomics data, protein identification, quantification results and statistical analysis are available upon request and will be fully available upon publication. There are no restrictions on materials other than those imposed by the commercial availability of cell lines, antibodies, drugs, and other reagents used in this study.

## References and Notes

1. S. Klaeger et al., The target landscape of clinical kinase drugs. Science 358 (2017).

2. M. Frejno et al., Proteome activity landscapes of tumor cell lines determine drug responses. Nature Communications 2020 11:1 11, 1–12 (2020).

3. A. Lin et al., Off-target toxicity is a common mechanism of action of cancer drugs undergoing clinical trials. Science Translational Medicine 11, 8412 (2019).

4. M. Niepel et al., Common and cell-type specific responses to anti-cancer drugs revealed by high throughput transcript profiling. Nature Communications 2017 8:1 8, 1–11 (2017).

5. J. Zecha et al., Decrypting drug actions and protein modifications by dose- and time-resolved proteomics. Science 380, 93–101 (2023).

6. Y. C. Chang et al., Decrypting lysine deacetylase inhibitor action and protein modifications by dose-resolved proteomics. Cell Reports 43, 114272 (2024).

7. L. Litichevskiy et al., A Library of Phosphoproteomic and Chromatin Signatures for Characterizing Cellular Responses to Drug Perturbations. Cell Systems 6, 424–443.e7 (2018).

8. E. B. Ünal et al., A compendium of ERK targets. FEBS Letters 591, 2607–2615 (2017).

9. H. S. Solanki et al., Cell-type Specific Adaptive Signaling Responses to KRASG12C inhibition. Clinical cancer research: an official journal of the American Association for Cancer Research 27, 2533 (2021).

10. S. A. Stuart et al., A Phosphoproteomic Comparison of B-RAF V600E and MKK1/2 Inhibitors in Melanoma Cells* □ S. Molecular & Cellular Proteomics 14, 1599–1615 (2015).

11. J. E. Klomp et al., Determining the ERK-regulated phosphoproteome driving KRAS-mutant cancer. Science (New York, N.Y.) 384, eadk0850 (2024).

12. L. Huang et al., KRAS mutation: from undruggable to druggable in cancer. Signal Transduction and Targeted Therapy 2021 6:1 6, 1–20 (2021).

13. B. A. Lanman et al., Discovery of a Covalent Inhibitor of KRASG12C (AMG 510) for the Treatment of Solid Tumors. Journal of Medicinal Chemistry 63, 52–65 (2020).

14. J. B. Fell et al., Identification of the Clinical Development Candidate MRTX849, a Covalent KRASG12CInhibitor for the Treatment of Cancer. Journal of Medicinal Chemistry 63, 6679–6693 (2020).

15. J. Popow et al., Targeting cancer with small-molecule pan-KRAS degraders. Science 385, 1338–1347 (2024).

16. D. Kim et al., Pan-KRAS inhibitor disables oncogenic signalling and tumour growth. Nature 2023 619:7968 619, 160–166 (2023).

17. A. R. Moore et al., RAS-targeted therapies: is the undruggable drugged? Nature Reviews Drug Discovery 2020 19:8 19, 533–552 (2020).

18. M. Holderfield et al., Concurrent inhibition of oncogenic and wild-type RAS-GTP for cancer therapy. Nature 2024 629:8013 629, 919–926 (2024).

19. L. Lautenbacher et al., ProteomicsDB: toward a FAIR open-source resource for life-science research. Nucleic Acids Research 50, D1541–D1552 (2022).

20. S. Eckert et al., Decrypting the molecular basis of cellular drug phenotypes by dose-resolved expression proteomics. Nature Biotechnology 2024, 1–10 (2024).

21. M. Kuljanin et al., Reimagining high-throughput profiling of reactive cysteines for cell-based screening of large electrophile libraries. Nature Biotechnology 2021 39:5 39, 630–641 (2021).

22. F. P. Bayer et al., CurveCurator: a recalibrated F-statistic to assess, classify, and explore significance of dose–response curves. Nature Communications 14 (2023).

23. J. L. Johnson et al., An atlas of substrate specificities for the human serine/threonine kinome. Nature 613, 759–766 (2023).

24. T. M. Yaron-Barir et al., The intrinsic substrate specificity of the human tyrosine kinome. Nature 2024 629:8014 629, 1174–1181 (2024).

25. R. Anjum et al., The RSK family of kinases: emerging roles in cellular signalling. Nature Reviews Molecular Cell Biology 2008 9:10 9, 747–758 (2008).

26. D. N. Sgouras et al., ERF: an ETS domain protein with strong transcriptional repressor activity, can suppress ets associated tumorigenesis and is regulated by phosphorylation during cell cycle and mitogenic stimulation. The EMBO Journal 14, 4781–4793 (1995).

27. J. M. Bomberger et al., The deubiquitinating enzyme USP10 regulates the post-endocytic sorting of cystic fibrosis transmembrane conductance regulator in airway epithelial cells. The Journal of biological chemistry 284, 18778–18789 (2009).

28. J. Yuan et al., USP10 regulates p53 localization and stability by deubiquitinating p53. Cell 140, 384–396 (2010).

29. M. Courcelles et al., Phosphoproteome dynamics reveal novel ERK1/2 MAP kinase substrates with broad spectrum of functions. Molecular Systems Biology 9, 669 (2013).

30. C. Pan et al., Global Effects of Kinase Inhibitors on Signaling Networks Revealed by Quantitative Phosphoproteomics. Molecular & Cellular Proteomics: MCP 8, 2796 (2009).

31. M. B. Ryan et al., Vertical pathway inhibition overcomes adaptive feedback resistance to KRASG12C inhibition. Clinical cancer research: an official journal of the American Association for Cancer Research 26, 1633 (2020).

32. J. Y. Xue et al., Rapid non-uniform adaptation to conformation-specific KRASG12C inhibition. Nature 577, 421 (2020).

33. C. H. Chen et al., MEK inhibitors induce Akt activation and drug resistance by suppressing negative feedback ERK mediated HER2 phosphorylation at Thr701. Molecular Oncology 11, 1273 (2017).

34. R. G. Hodge et al., Regulating Rho GTPases and their regulators. Nature Reviews Molecular Cell Biology 2016 17:8 17, 496–510 (2016).

35. K. Stamatiou et al., CCDC86 is a novel Ki-67-interacting protein important for cell division. Journal of Cell Science 136 (2023).

36. J. A. Klomp et al., Defining the KRAS- and ERK-dependent transcriptome in KRAS-mutant cancers. Science (New York, N.Y.) 384, eadk0775 (2024).

37. Z. S. Hann et al., Structural basis for adenylation and thioester bond formation in the ubiquitin E1. Proceedings of the National Academy of Sciences of the United States of America 116, 15475–15484 (2019).

38. A. K. L. Liess et al., Autoinhibition Mechanism of the Ubiquitin-Conjugating Enzyme UBE2S by Autoubiquitination. Structure 27, 1195–1210.e7 (2019).

39. G. Prus et al., Global, site-resolved analysis of ubiquitylation occupancy and turnover rate reveals systems properties. doi: 10.1016/j.cell.2024.03.024 (2024).

40. H. Miyashita et al., KRAS G12C inhibitor combination therapies: current evidence and challenge. Frontiers in Oncology 14, 1380584 (2024).

41. K. Fukuda et al., Targeting WEE1 enhances the antitumor effect of KRAS-mutated non-small cell lung cancer harboring TP53 mutations. Cell Reports Medicine 5 (2024).

42. J. Zecha et al., TMT Labeling for the Masses: A Robust and Cost-efficient, In-solution Labeling Approach. Molecular & Cellular Proteomics 18, 1468–1478 (2019).

43. J. Cox et al., MaxQuant enables high peptide identification rates, individualized p.p.b.-range mass accuracies and proteome-wide protein quantification. Nature Biotechnology 2008 26:12 26, 1367–1372 (2008).

44. J. Cox et al., Andromeda: A peptide search engine integrated into the MaxQuant environment. Journal of Proteome Research 10, 1794–1805 (2011).

45. V. Sharma et al., Panorama Public: A Public Repository for Quantitative Data Sets Processed in Skyline. Molecular & Cellular Proteomics 17, 1239–1244 (2018).

46. J. Jumper et al., Highly accurate protein structure prediction with AlphaFold. Nature 2021 596:7873 596, 583–589 (2021).

47. Z. Lv et al., Crystal structure of a human ubiquitin E1–ubiquitin complex reveals conserved functional elements essential for activity. The Journal of Biological Chemistry 293, 18337 (2018).

48. F. Hamood et al., SIMSI-Transfer: Software-Assisted Reduction of Missing Values in Phosphoproteomic and Proteomic Isobaric Labeling Data Using Tandem Mass Spectrum Clustering. Molecular & Cellular Proteomics 21, 100238 (2022).

49. S. Berg et al., ilastik: interactive machine learning for (bio)image analysis. Nature Methods 2019 16:12 16, 1226–1232 (2019).

50. P. Vagnarelli et al., Repo-Man Coordinates Chromosomal Reorganization with Nuclear Envelope Reassembly during Mitotic Exit. Developmental Cell 21, 328–342 (2011).

51. M. Gemmer et al., Visualization of translation and protein biogenesis at the ER membrane. Nature 2023 614:7946 614, 160–167 (2023).

52. S. Riniker et al., Better Informed Distance Geometry: Using What We Know to Improve Conformation Generation. Journal of Chemical Information and Modeling 55, 2562–2574 (2015).

53. B. Ruprecht et al., High pH Reversed-Phase Micro-Columns for Simple, Sensitive, and Efficient Fractionation of Proteome and (TMT labeled) Phosphoproteome Digests. Methods in molecular biology (Clifton, N.J.) 1550, 83–98 (2017).

54. N. D. Udeshi et al., Rapid and deep-scale ubiquitylation profiling for biology and translational research. Nature Communications 2020 11:1 11, 1–11 (2020).

55. M. Reinecke et al., Chemoproteomic Selectivity Profiling of PIKK and PI3K Kinase Inhibitors. ACS Chemical Biology 14, 655–664 (2019).

56. B. MacLean et al., Skyline: an open source document editor for creating and analyzing targeted proteomics experiments. Bioinformatics 26, 966–968 (2010).

57. J. Cox et al., Accurate Proteome-wide Label-free Quantification by Delayed Normalization and Maximal Peptide Ratio Extraction, Termed MaxLFQ. Molecular & Cellular Proteomics: MCP 13, 2513 (2014).

58. C. Ritz et al., Dose-Response Analysis Using R. PLOS ONE 10, e0146021 (2015).

59. N. T. Doncheva et al., Cytoscape StringApp: Network Analysis and Visualization of Proteomics Data. Journal of Proteome Research 18, 623–632 (2019).

60. J. Müller, et al., PTMNavigator: Interactive Visualization of Differentially Regulated Post-Translational Modifications in Cellular Signaling Pathways. bioRxiv, 2023.08.31.555601 (2024).

61. A. Bateman et al., UniProt: the universal protein knowledgebase in 2021. Nucleic Acids Research 49, D480–D489 (2021).

62. S. A. Lambert et al., The Human Transcription Factors. Cell 172, 650–665 (2018).

63. Z. J. Zhao et al., Purification and Cloning of PZR, a Binding Protein and Putative Physiological Substrate of Tyrosine Phosphatase SHP-2. Journal of Biological Chemistry 273, 29367–29372 (1998).

64. D. Lake et al., Negative feedback regulation of the ERK1/2 MAPK pathway. Cellular and Molecular Life Sciences: CMLS 73, 4397 (2016).

