## Supplementary Figures for "Illuminating oncogenic KRAS signaling by multi-dimensional chemical proteomics"

**This PDF file includes:**

Figs. S1 to S18

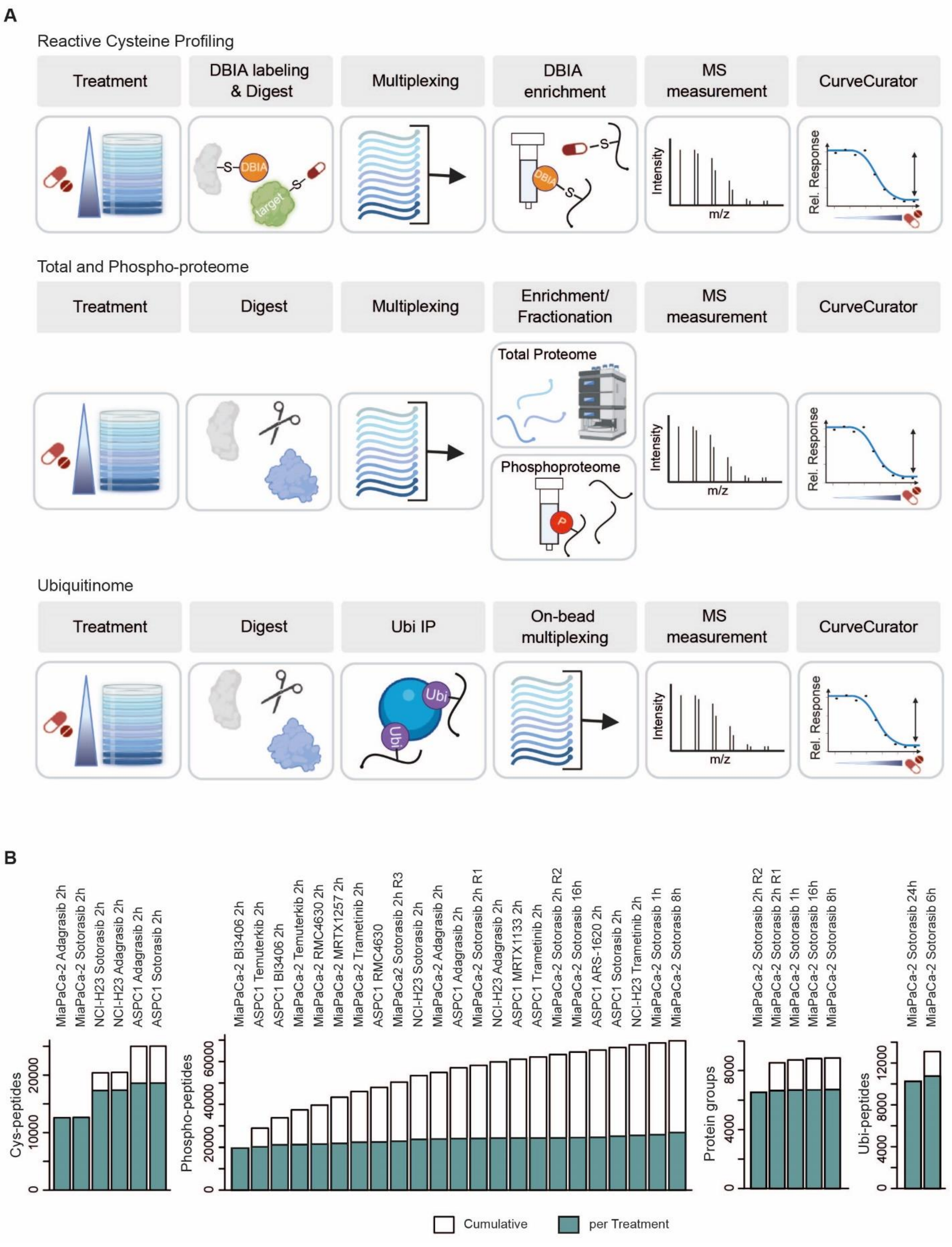
A
Reactive Cysteine Profiling
Treatment
DBIA labeling & Digest
Multiplexing
DBIA enrichment
MS measurement
CurveCurator
DBIA
target
Intensity
m/z
Rel. Response
Total and Phospho-proteome
Treatment
Digest
Multiplexing
Enrichment/ Fractionation
MS measurement
CurveCurator
Total Proteome
Phosphoproteome
P
Ubiquitinome
Treatment
Digest
Ubi IP
On-bead multiplexing
MS measurement
CurveCurator
Ubi
B
MiaPaCa-2 Adagrasib 2h
MiaPaCa-2 Sotorasib 2h
NCI-H23 Sotorasib 2h
NCI-H23 Adagrasib 2h
ASPC1 Adagrasib 2h
ASPC1 Sotorasib 2h
MiaPaCa-2 BI3406 2h
ASPC1 Temuterkib 2h
ASPC1 BI3406 2h
MiaPaCa-2 Temuterkib 2h
MiaPaCa-2 RMC4630 2h
MiaPaCa-2 MRTX1257 2h
MiaPaCa-2 Trametinib 2h
ASPC1 RMC4630
MiaPaCa2 Sotorasib 2h R3
NCI-H23 Sotorasib 2h
MiaPaCa-2 Adagrasib 2h
ASPC1 Adagrasib 2h
MiaPaCa-2 Sotorasib 2h R1
NCI-H23 Adagrasib 2h
ASPC1 MRTX1133 2h
ASPC1 Trametinib 2h
MiaPaCa-2 Sotorasib 2h R2
MiaPaCa-2 Sotorasib 16h
ASPC1 ARS-1620 2h
ASPC1 Sotorasib 2h
NCI-H23 Trametinib 2h
MiaPaCa-2 Sotorasib 1h
MiaPaCa-2 Sotorasib 8h
MiaPaCa-2 Sotorasib 2h R2
MiaPaCa-2 Sotorasib 2h R1
MiaPaCa-2 Sotorasib 1h
MiaPaCa-2 Sotorasib 16h
MiaPaCa-2 Sotorasib 8h
MiaPaCa-2 Sotorasib 24h
MiaPaCa-2 Sotorasib 6h
Cys-peptides
0
10000
20000
Phospho-peptides
0
20000
40000
60000
Protein groups
0
4000
8000
Ubi-peptides
0
4000
8000
12000
Cumulative
per Treatment

**Fig. S1: Chemical proteomics workflows applied in this study.**

**(A)** Upper Panel: For reactive cysteine profiling, cells were treated with increasing concentrations of a drug. Following cell lysis, reactive cysteines were labeled with desthiobiotin-iodoacetamide (DBIA). Proteins were then digested with trypsin, followed by stable isotope labeling using tandem mass tags (TMT; one tag for each drug concentration), combining samples (multiplexing) and Streptavidin enrichment of cysteine-containing peptides (cys-peptides). Sample were analyzed by LC-MS3, and the resulting quantitative data were fitted by CurveCurator using a sigmoidal regression model to generate dose-response curves for each peptide. Statistical assessment was performed by CurveCurator. Middle Panel: Full (total) proteomes and phosphoproteomes were processed as above but without DBIA labeling. Enrichment of phosphorylated peptides (phospho-peptides) was performed using immobilized metal affinity chromatography (IMAC). Bottom Panel: The ubiquitinome was isolated as above but with the following modifications. Enrichment of ubiquitinated peptides (ubi-peptides) was achieved by immunoprecipitation using antibody-coupled beads that bind to the Gly-Gly remnant of ubiquitin generated by tryptic digestion. Bead-bound peptides were labeled with TMT while still on the beads, and the eluate was processed as above. See materials and methods for details.

**(B)** Summary of all data sets in this study. Barplots showing the number of identified cys-peptides (left barplot), phospho-peptides (second left), protein groups (second right) and ubi-peptides (right barplot). The blue parts of each bar represents the number of identifications for a given experiment. The grey parts show the progressive accumulation of unique (non-redundant) peptides/protein groups across experiments.

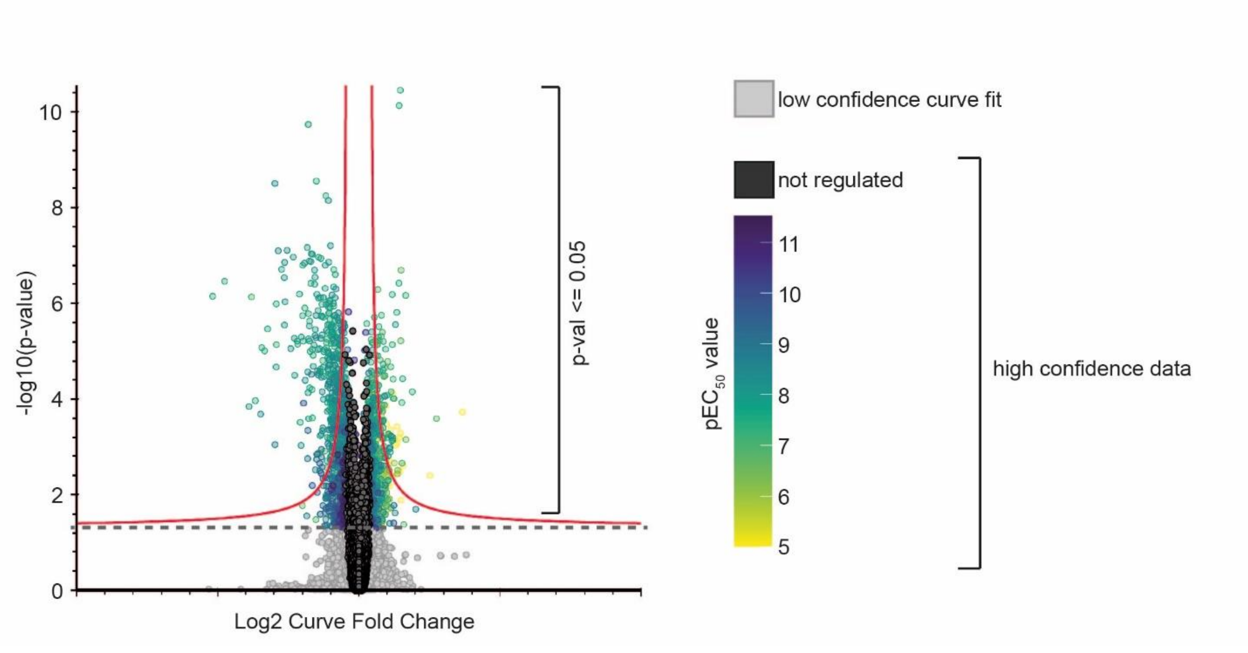


**Fig. S2 Statistical evaluation of dose-response data using CurveCurator.**

**(A)** Example volcano plot showing results of CurveCurator processing. Throughout the manuscript, we refer to statistically significantly regulated data if e. g. phospho-peptides are outside the red asymptote according to F-statistic (cutoff values: fold change-value = 0.6 for cysteine profiling data and 0.45 for all other data; p-value = 0.05). CurveCurator output data was further filtered by additional criteria to remove data influenced only by the first or last data point (see materials and methods for details, Classification). All downstream analysis and comparisons (using curve fold change or $pEC_{50}$ dimensions) were performed on high confidence data only. $pEC_{50}$ values were only used for phospho-peptides that were clearly classified as either "up" or "down" regulated according to all applied criteria.

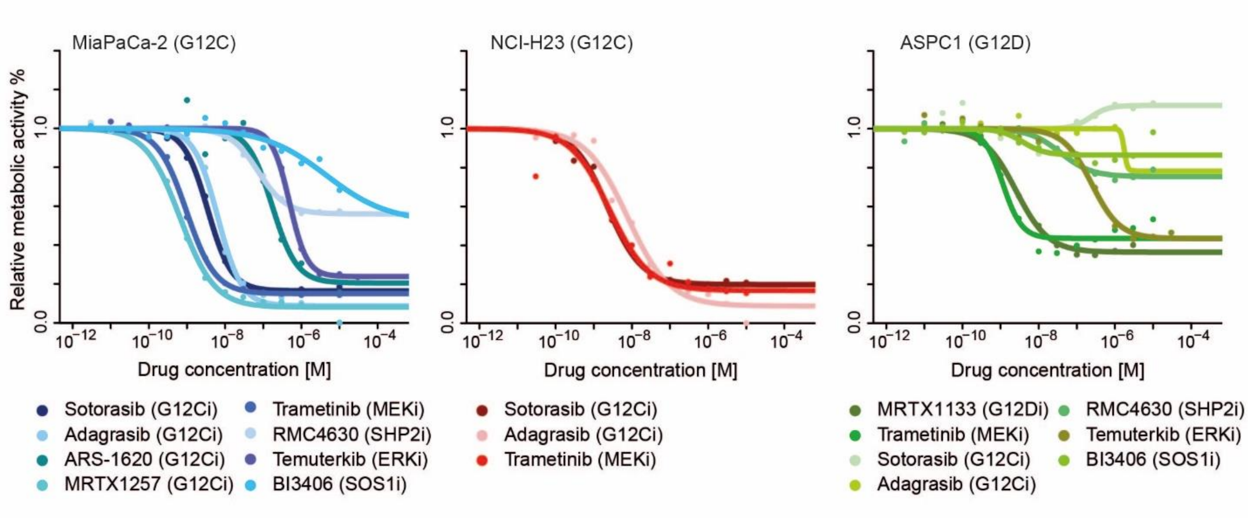


**Fig. S3: Cell viability analysis in response to inhibitors.**
MiaPaCa-2 (left), NCI-H23 (middle) and ASPC1 (right) cells were treated with increasing concentrations of drugs for 72h and cell viability was determined using a metabolic readout (Alamar blue assay).

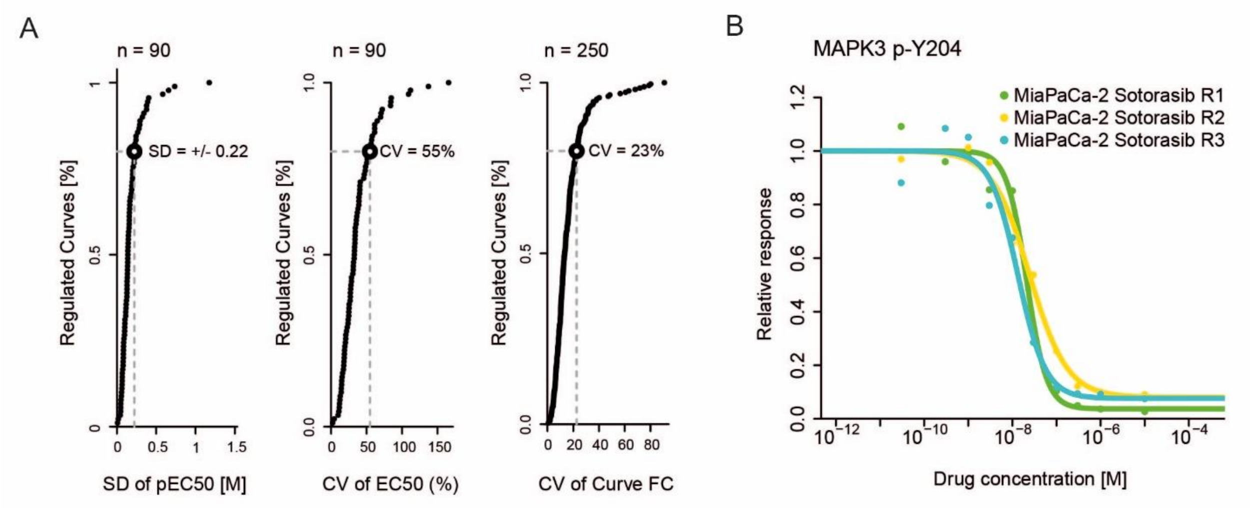


**Fig. S4: Reproducibility analysis of decryptM data.**
**(A)** Cumulative density plots of drug-regulated phospho-peptide dose-response curves as a function of: i) the coefficient of variation of $pEC_{50}$ ($-logEC_{50}$) values (phospho-peptides regulated across all replicates) (left), ii) standard deviation of $EC_{50}$ values (middle; phospho-peptides regulated across all replicates) and CV of response (right) from replicate analysis (n=3, biological replicates) of MiaPaCa-2 cells treated with Sotorasib (2h). CV of response was based on curve fold change of regulated phospho-peptides in at least one replicate plus curve fold change of the same phospho-peptides with high-confidence curve-fit from other replicates; see fig. S2A). Numbers at the top of each plot denotes the number of phospho-peptides that form the basis for the analysis.
**(B)** Example replicate dose-response curves from (A).

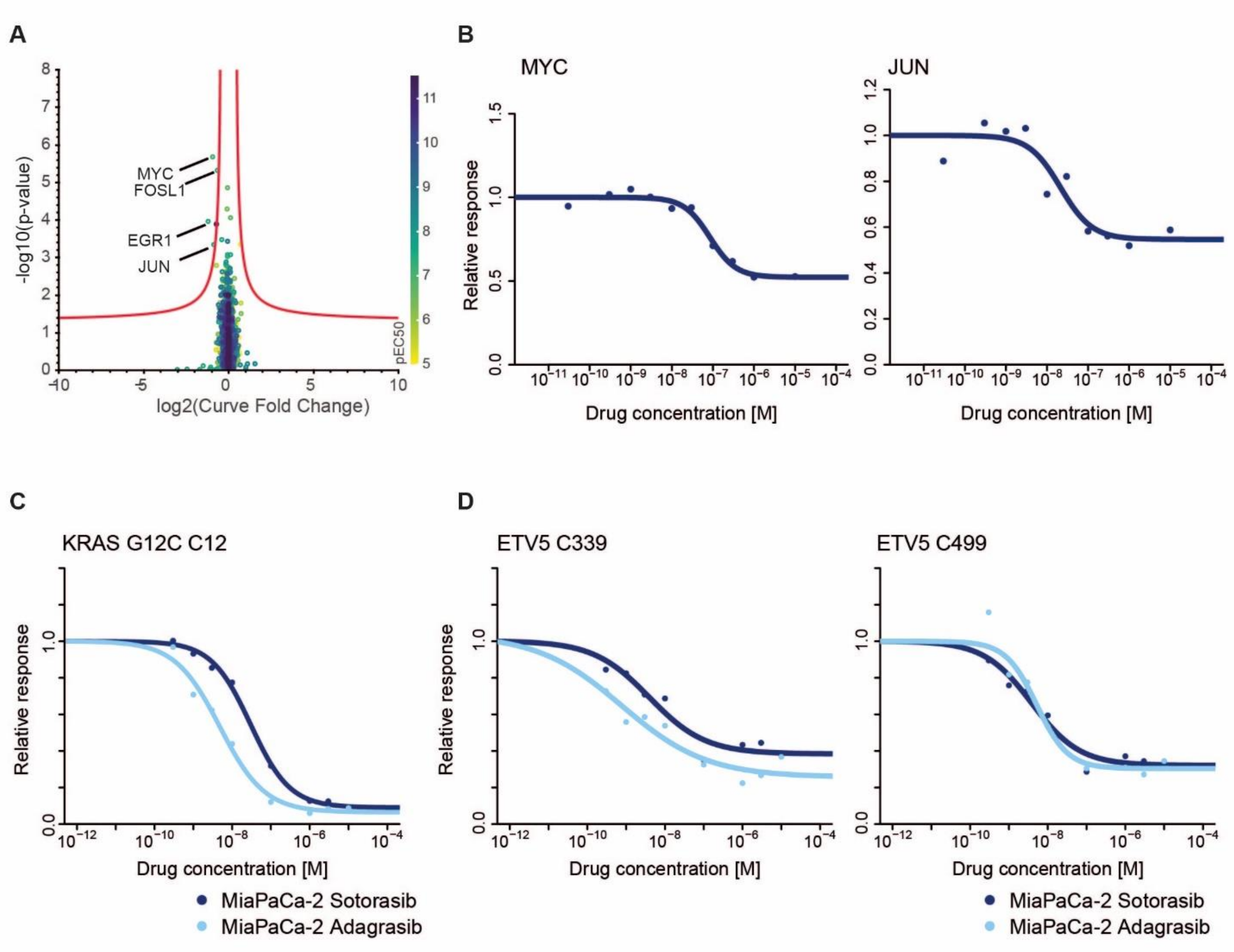


**Fig. S5: Drug-induced protein abundance changes.**

**(A)** Volcano plot from CurveCurator analysis of dose-dependent protein abundance changes in MiaPaCa-2 cells treated with Sotorasib (2h).

**(B)** Examples for dose-response curves of protein abundance changes of MYC and JUN from panel (A).

**(C)** Dose-response curves of reactive cysteine profiling experiments for KRAS G12C in MiaPaCa-2 (left) and two different ETV5 cys-peptides (middle and right) in response to 2-hour Sotorasib and Adagrasib treatment. The near identical characteristics of both ETV5 sites to that of KRAS G12C in response to both drugs most likely means that these apparent changes result from changes in protein expression and not from covalent drug binding (as observed for the proteins shown in panels (A) and (B); ETV5 was only detected in reactive cysteine profiling but not in full proteome profiling experiments).

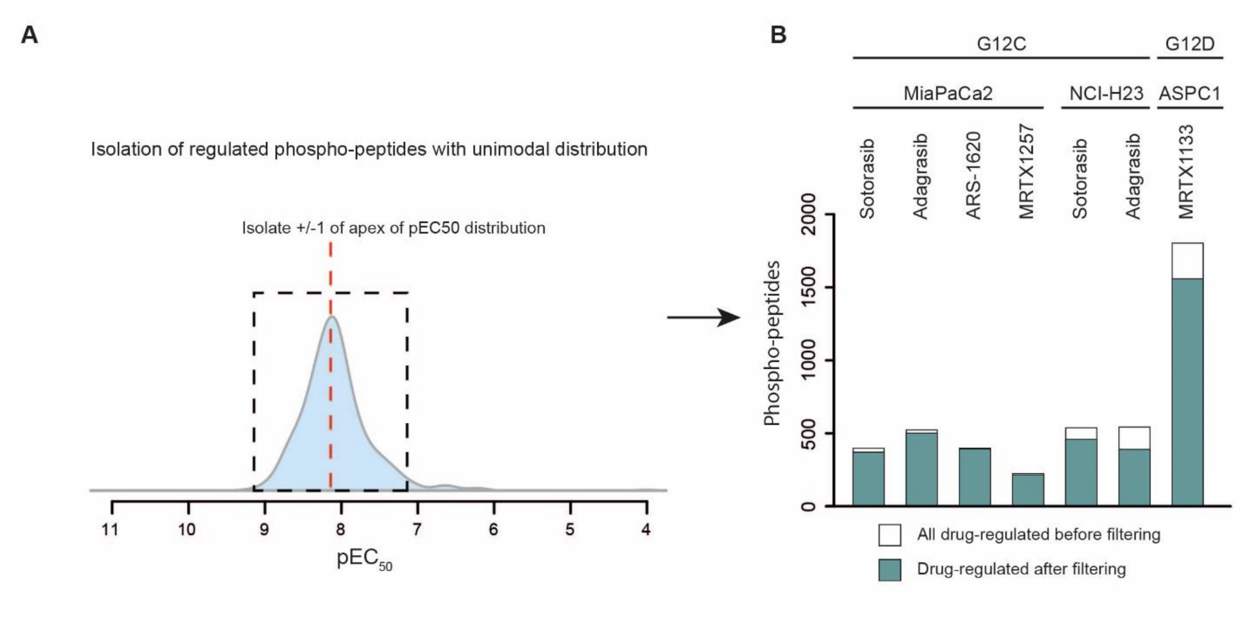


**Fig. S6: KRAS-dependent regulated phospho-peptides.**
**(A)** Schematic representation of the filtering approach taken for KRAS inhibiton-induced phospho-peptide abundance changes. Briefly, the apex of the $pEC_{50}$ distribution of all drug-regulated phospho-peptides was determined for each KRASi treatment. Only phospho-peptides with $pEC_{50}$ values ± 1 around the apex were used for further analysis. The rationale for this filtering step is that phospho-peptides regulated by the same cellular mechanism should show drug-regulation at similar concentrations.
**(B)** Barplot showing the number of regulated phospho-peptides before (white) and after (green) filtering.

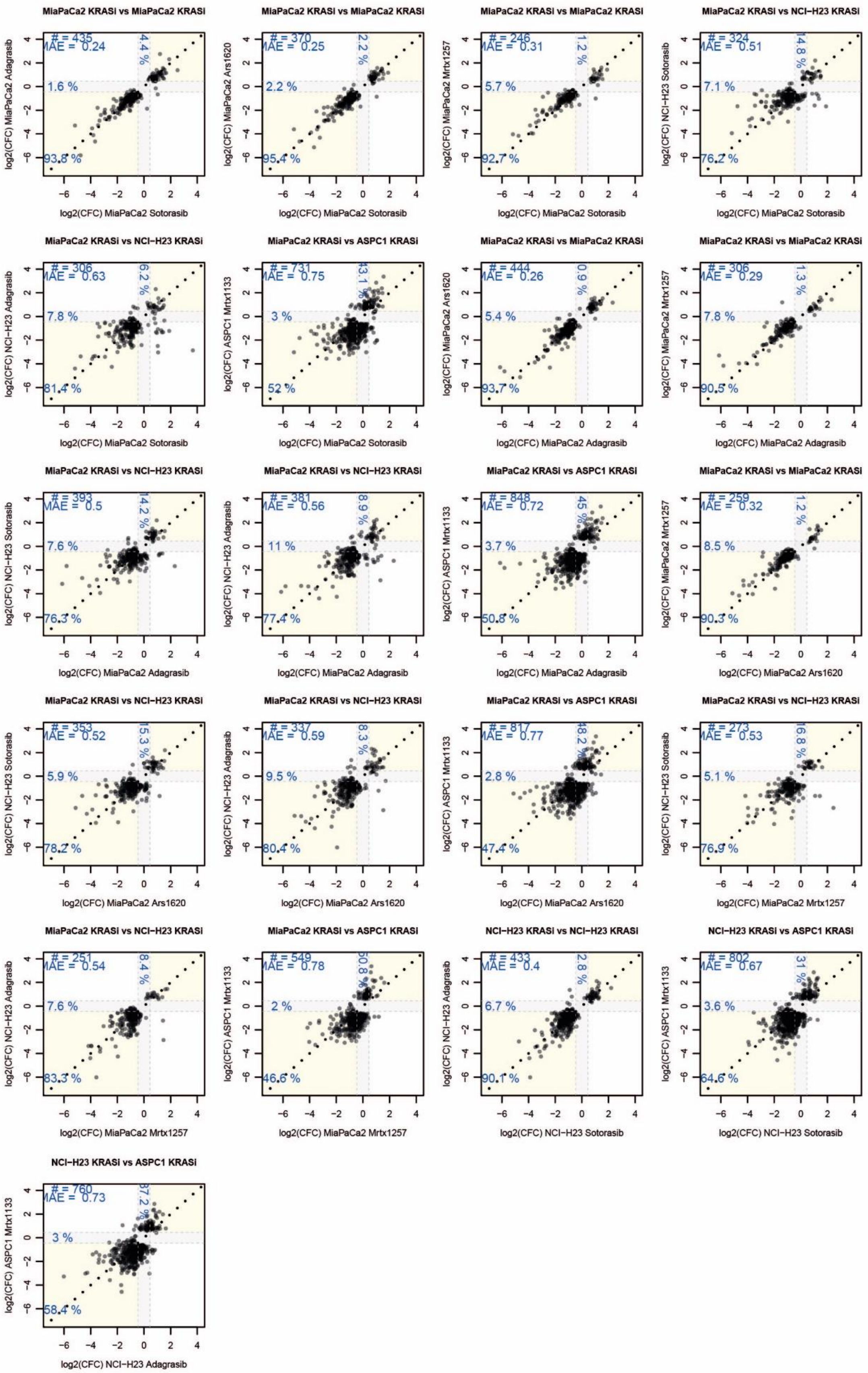
MiaPaCa2 KRASi vs MiaPaCa2 KRASi
# = 435
MAE = 0.24
4.4 %
1.6 %
93.8 %
log2(CFC) MiaPaCa2 Adagrasib
log2(CFC) MiaPaCa2 Sotorasib
MiaPaCa2 KRASi vs MiaPaCa2 KRASi
# = 370
MAE = 0.25
2.2 %
2.2 %
95.4 %
log2(CFC) MiaPaCa2 Ars1620
log2(CFC) MiaPaCa2 Sotorasib
MiaPaCa2 KRASi vs MiaPaCa2 KRASi
# = 246
MAE = 0.31
1.2 %
5.7 %
92.7 %
log2(CFC) MiaPaCa2 Mrtx1257
log2(CFC) MiaPaCa2 Sotorasib
MiaPaCa2 KRASi vs NCI−H23 KRASi
# = 324
MAE = 0.51
14.8 %
7.1 %
76.2 %
log2(CFC) NCI−H23 Sotorasib
log2(CFC) MiaPaCa2 Sotorasib
MiaPaCa2 KRASi vs NCI−H23 KRASi
# = 306
MAE = 0.63
6.2 %
7.8 %
81.4 %
log2(CFC) NCI−H23 Adagrasib
log2(CFC) MiaPaCa2 Sotorasib
MiaPaCa2 KRASi vs ASPC1 KRASi
# = 731
MAE = 0.75
43.1 %
3 %
52 %
log2(CFC) ASPC1 Mrtx1133
log2(CFC) MiaPaCa2 Sotorasib
MiaPaCa2 KRASi vs MiaPaCa2 KRASi
# = 444
MAE = 0.26
0.9 %
5.4 %
93.7 %
log2(CFC) MiaPaCa2 Ars1620
log2(CFC) MiaPaCa2 Adagrasib
MiaPaCa2 KRASi vs MiaPaCa2 KRASi
# = 306
MAE = 0.29
1.3 %
7.8 %
90.5 %
log2(CFC) MiaPaCa2 Mrtx1257
log2(CFC) MiaPaCa2 Adagrasib
MiaPaCa2 KRASi vs NCI−H23 KRASi
# = 393
MAE = 0.5
14.2 %
7.6 %
76.3 %
log2(CFC) NCI−H23 Sotorasib
log2(CFC) MiaPaCa2 Adagrasib
MiaPaCa2 KRASi vs NCI−H23 KRASi
# = 381
MAE = 0.56
8.9 %
11 %
77.4 %
log2(CFC) NCI−H23 Adagrasib
log2(CFC) MiaPaCa2 Adagrasib
MiaPaCa2 KRASi vs ASPC1 KRASi
# = 848
MAE = 0.72
45 %
3.7 %
50.8 %
log2(CFC) ASPC1 Mrtx1133
log2(CFC) MiaPaCa2 Adagrasib
MiaPaCa2 KRASi vs MiaPaCa2 KRASi
# = 259
MAE = 0.32
1.2 %
8.5 %
90.3 %
log2(CFC) MiaPaCa2 Mrtx1257
log2(CFC) MiaPaCa2 Ars1620
MiaPaCa2 KRASi vs NCI−H23 KRASi
# = 353
MAE = 0.52
15.3 %
5.9 %
78.2 %
log2(CFC) NCI−H23 Sotorasib
log2(CFC) MiaPaCa2 Ars1620
MiaPaCa2 KRASi vs NCI−H23 KRASi
# = 337
MAE = 0.59
8.3 %
9.5 %
80.4 %
log2(CFC) NCI−H23 Adagrasib
log2(CFC) MiaPaCa2 Ars1620
MiaPaCa2 KRASi vs ASPC1 KRASi
# = 817
MAE = 0.77
48.2 %
2.8 %
47.4 %
log2(CFC) ASPC1 Mrtx1133
log2(CFC) MiaPaCa2 Ars1620
MiaPaCa2 KRASi vs NCI−H23 KRASi
# = 273
MAE = 0.53
16.8 %
5.1 %
76.9 %
log2(CFC) NCI−H23 Sotorasib
log2(CFC) MiaPaCa2 Mrtx1257
MiaPaCa2 KRASi vs NCI−H23 KRASi
# = 251
MAE = 0.54
8.4 %
7.6 %
83.3 %
log2(CFC) NCI−H23 Adagrasib
log2(CFC) MiaPaCa2 Mrtx1257
MiaPaCa2 KRASi vs ASPC1 KRASi
# = 549
MAE = 0.78
50.8 %
2 %
46.6 %
log2(CFC) ASPC1 Mrtx1133
log2(CFC) MiaPaCa2 Mrtx1257
NCI−H23 KRASi vs NCI−H23 KRASi
# = 433
MAE = 0.4
2.8 %
6.7 %
90.1 %
log2(CFC) NCI−H23 Adagrasib
log2(CFC) NCI−H23 Sotorasib
NCI−H23 KRASi vs ASPC1 KRASi
# = 802
MAE = 0.67
31 %
3.6 %
64.6 %
log2(CFC) ASPC1 Mrtx1133
log2(CFC) NCI−H23 Sotorasib
NCI−H23 KRASi vs ASPC1 KRASi
# = 760
MAE = 0.73
37.2 %
3 %
58.4 %
log2(CFC) ASPC1 Mrtx1133
log2(CFC) NCI−H23 Adagrasib

**Fig. S7: Comparison of responses to different KRAS inhibitors.**

Scatter plots showing the pairwise comparison of regulated phospho-peptide responses for any combination of KRAS inhibitor and cell line. MAE: mean absolute error of log2 curve fold changes; # number of phospho-peptides in plot; Dotted lines mark the boundaries of CurveCurator log2 fold change cut-off (fc-values = ±0.45). Yellow areas contain phospho-peptides with consistent responses, grey areas contain phospho-peptides regulated by one of the conditions only. Percentages denote the relative proportion of phospho-peptides in the yellow and grey areas respectively. Values on the y- and x-axis are the log2 curve fold change (CFC) of phospho-peptides in response to drug.

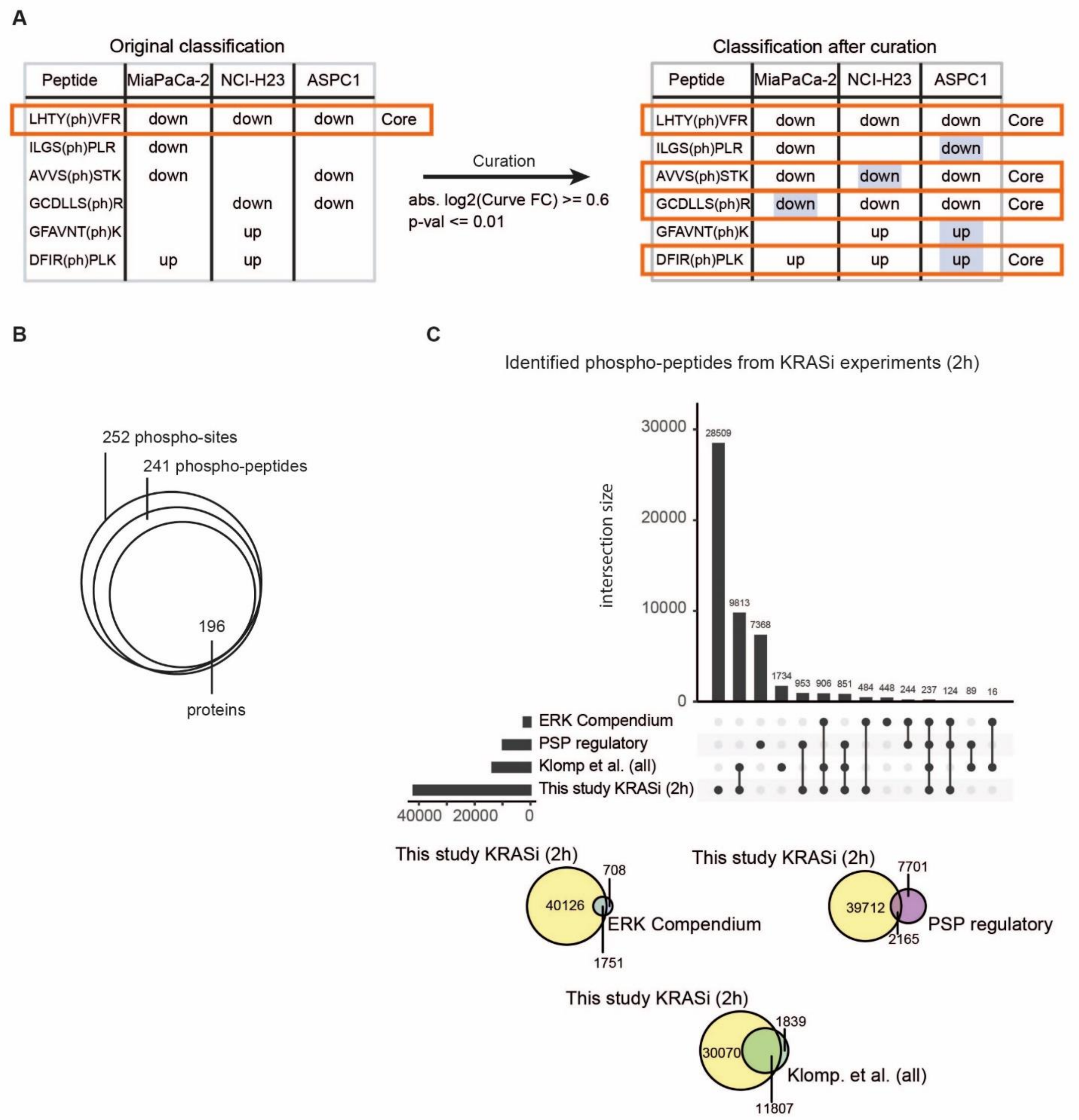


**Fig. S8: KRAS signaling signature and data comparison**

**(A)** Exemplification of the curation process for selecting phospho-peptides for inclusion in the KRAS core signaling signature (see extended methods for details). Briefly, phospho-peptides meeting CurveCurator criteria for drug regulation by at least one KRAS drug in all three cell lines were included in the signature (left panel). Phospho-peptides that met CurveCurator criteria in at least one but not all cell lines (because the required fold change response was not quite reached in a particular cell line) were also included as long as they showed a CurveCurator p-value of $\leq 0.01$, an absolute log2 fold change (response) of $\geq 0.6$ (right panel) and followed the same quality criteria as regular curves ($EC_{50}$ must lie within the treatment range, see materials and methods (classification) for details).

**(B)** Venn diagram of phospho-sites, phospho-peptides and proteins (based on UniProt canonical sequences of protein coding genes) that constitute the KRAS core signaling signature.

**(C)** Upper panel: comparison and overlap of all identified phospho-sites from experiments performed with KRAS inhibitors for 2h in this study, the ERK-dependent phospho-proteome (Klomp et al.; all experiments; (*11*)), compiled in the ERK compendium (*8*) and annotated as 'regulated' in Phosphositeplus.org (PSP regulatory). Only phospho-sites from canonical protein sequences were compared. Lower panel: Venn diagrams depicting the overlap of identified phospho-peptides from this study with those from other resources.

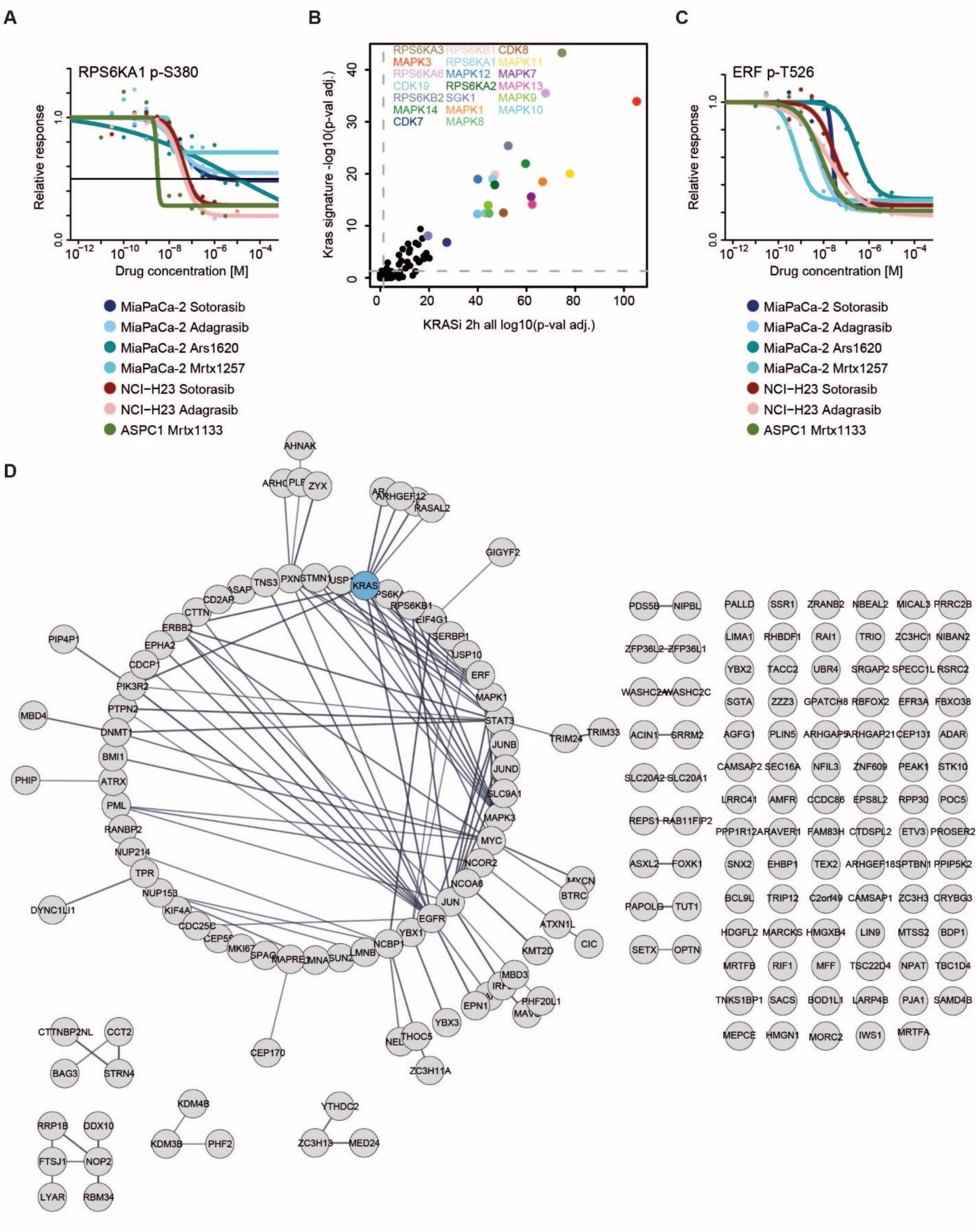
A
RPS6KA1 p-S380
Relative response
Drug concentration [M]
MiaPaCa-2 Sotorasib
MiaPaCa-2 Adagrasib
MiaPaCa-2 Ars1620
MiaPaCa-2 Mrtx1257
NCI−H23 Sotorasib
NCI−H23 Adagrasib
ASPC1 Mrtx1133
B
RPS6KA3 RPS6KB1 CDK8
MAPK3 RPS6KA1 MAPK11
RPS6KA6 MAPK12 MAPK7
CDK19 RPS6KA2 MAPK13
RPS6KB2 SGK1 MAPK9
MAPK14 MAPK1 MAPK10
CDK7 MAPK8
Kras signature -log10(p-val adj.)
KRASi 2h all log10(p-val adj.)
C
ERF p-T526
Relative response
Drug concentration [M]
MiaPaCa-2 Sotorasib
MiaPaCa-2 Adagrasib
MiaPaCa-2 Ars1620
MiaPaCa-2 Mrtx1257
NCI−H23 Sotorasib
NCI−H23 Adagrasib
ASPC1 Mrtx1133
D
AHNAK
ZYX
RASAL2
GIGYF2
KRAS
TNS3
PXN
STMN1
CD2AP
CTTN
ERBB2
EPHA2
CDCP1
PIK3R2
PTPN2
DNMT1
BMI1
ATRX
PML
RANBP2
NUP214
TPR
NUP153
KIF4A
CDC25C
MKI67
LMNB
NCBP1
YBX1
EGFR
JUN
NCOA6
NCOR2
MYC
MAPK3
SLC9A1
JUND
JUNB
STAT3
MAPK1
ERF
USP10
SERBP1
EIF4G1
RPS6KB1
PIP4P1
MBD4
PHIP
DYNC1LI1
TRIM24
TRIM33
BTRC
ATXN1L
CIC
KMT2D
MBD3
PHF20L1
EPN1
YBX3
THOC5
ZC3H11A
CEP170
CTTNBP2NL
CCT2
BAG3
STRN4
RRP1B
DDX10
FTSJ1
NOP2
LYAR
RBM34
KDM4B
KDM3B
PHF2
YTHDC2
ZC3H13
MED24
PDS5B
NIPBL
ZFP36L2
ZFP36L1
WASHC2A
WASHC2C
ACIN1
SRRM2
SLC20A2
SLC20A1
REPS1
RAB11FIP2
ASXL2
FOXK1
PAPOLG
TUT1
SETX
OPTN
PALLD
SSR1
ZRANB2
NBEAL2
MICAL3
PRRC2B
LIMA1
RHBDF1
RAI1
TRIO
ZC3HC1
NIBAN2
YBX2
TACC2
UBR4
SRGAP2
SPECC1L
RSRC2
SGTA
ZZZ3
GPATCH8
RBFOX2
EFR3A
FBXO38
AGFG1
PLIN5
ARHGAP21
CEP131
ADAR
CAMSAP2
SEC16A
NFIL3
ZNF609
PEAK1
STK10
LRRC41
AMFR
CCDC86
EPS8L2
RPP30
POC5
CTDSPL2
ETV3
PROSER2
SNX2
EHBP1
TEX2
ARHGEF18
PPIP5K2
BCL9L
TRIP12
C2orf49
CAMSAP1
ZC3H3
CRYBG3
HDGFL2
MARCKS
HMGXB4
LIN9
MTSS2
BDP1
MRTFB
RIF1
MFF
TSC22D4
NPAT
TBC1D4
TNKS1BP1
SACS
BOD1L1
LARP4B
PJA1
SAMD4B
MEPCE
HMGN1
MORC2
IWS1
MRTFA

**Fig. S9: Shared effects across cell lines treated with KRAS inhibitors.**
**(A)** Dose-response curves for the activity-determining phospho-site p-S380 of RPS6KA1, one of the most strongly enriched kinases in Figure 4C and panel (B).
**(B)** Scatter plot showing results of kinase motif enrichment analysis for phospho-peptides (down-regulated only) comprising the KRAS core signaling signature (y-axis) and all phospho-peptides down-regulated by KRAS inhibitors (x-axis).
**(C)** Dose-response curves for ERF T526 phosphorylation (known substrate of ERK) in response to different KRAS inhibitors in different cell lines.
**(D)** STRING network (confidence score ≥ 0.7 (high)) showing the interaction network of phospho-proteins underlying the phospho-peptides of the KRAS core signaling signature. KRAS itself was added manually as an additional node (cyan). The network shows that about half of the proteins of the core signaling signature had previously been functionally linked to each other, but the other half has not.

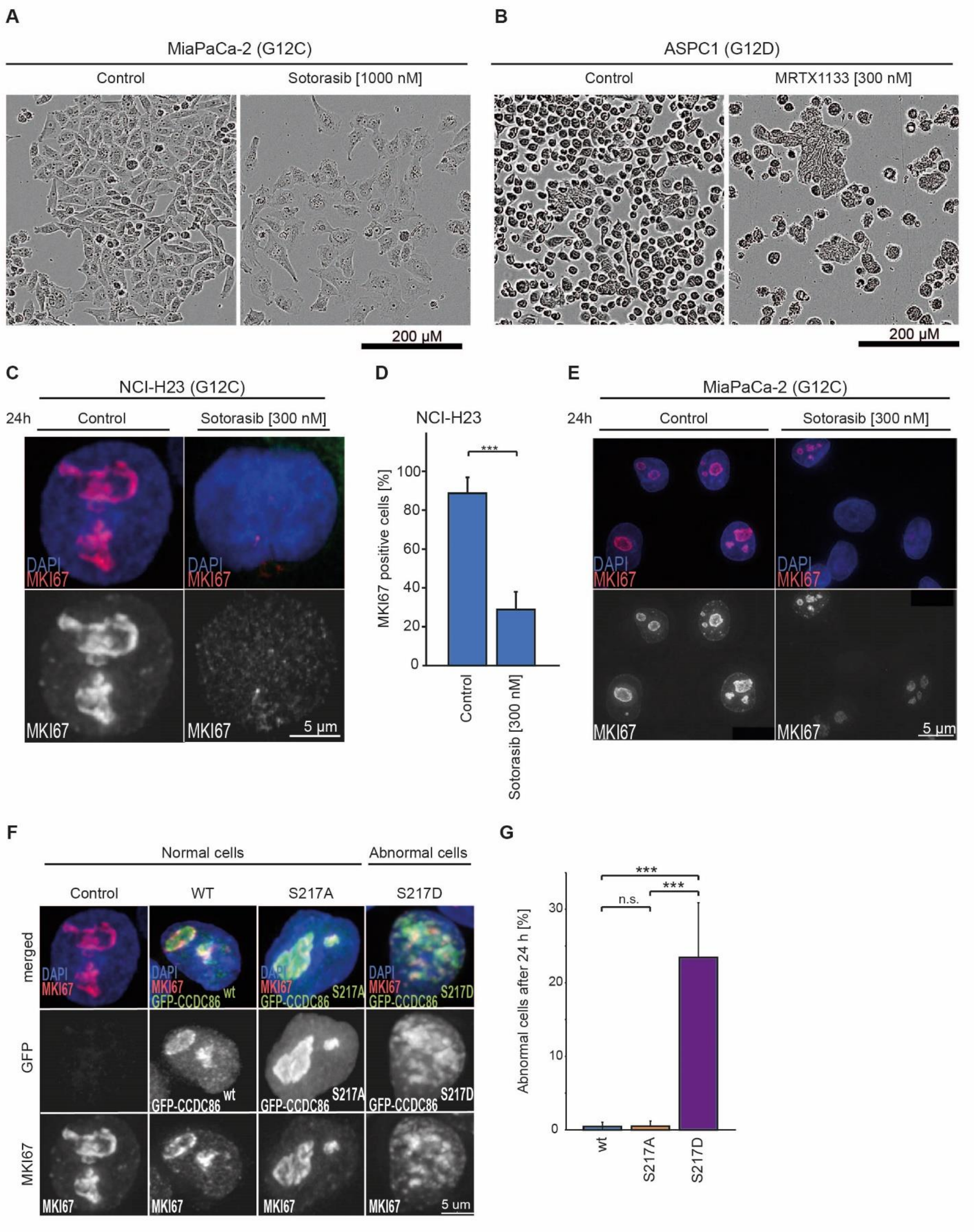


**Fig. S10: Phenotypic changes in response to KRAS Inhibition and CCDC86 variants.**

**(A)** Microscopic images illustrating morphological changes of MiaPaCa-2 cells in response to Sotorasib (72h). Scale bar 200 μm.
**(B)** same as panel (A) but for ASPC1 cells and Mrtx1133 treatment.
**(C)** Representative images of NCI-H23 cells in response to Sotorasib (24h). Cells were fixed and stained for MKI67 (red) and DNA (DAPI; blue). Scale bar 5 μm.
**(D)** Quantification of MKI67 positive cells from panel (C). The graph shows the average percentage of MKI67 positive cells of 4 biological replicates (DMSO; n=499. Sotorasib; n=489) and the error bars represent the standard deviations.
**(E)** Same as panel (C) but for MiaPaCa-2 cells.
**(F)** Representative images of NCI-H23 cells untransfected (control) or transfected with either GFP–CCDC86 WT, GFP–CCDC86 S217A or GFP–CCDC86 S217D constructs (green) for 24h. Cells were also stained for MKI67 (red) and DNA (DAPI). Scale bar 5 µm.
**(G)** Quantification of the cell phenotype from the experiment in Fig. 4G/ fig. S10F with abnormal MKI67 signal distribution as shown in (F) 24h post-transfection with GFP-CCDC86 WT, GFP-CCDC86 S217A and GFP-CCDC86 S217D. The graph shows the average percentage of abnormal MKI67 distribution cells of 2 biological replicates (GFP–CCDC86 WT; n=205. GFP–CCDC86 S217A; n=207. CCDC86 S217D; n=206) and the error bars represent the standard deviation. The data were statistically analyzed with a Chi-squared test. n.s., not significant; ***= $p < 0.001$, control: 0.1% DMSO.

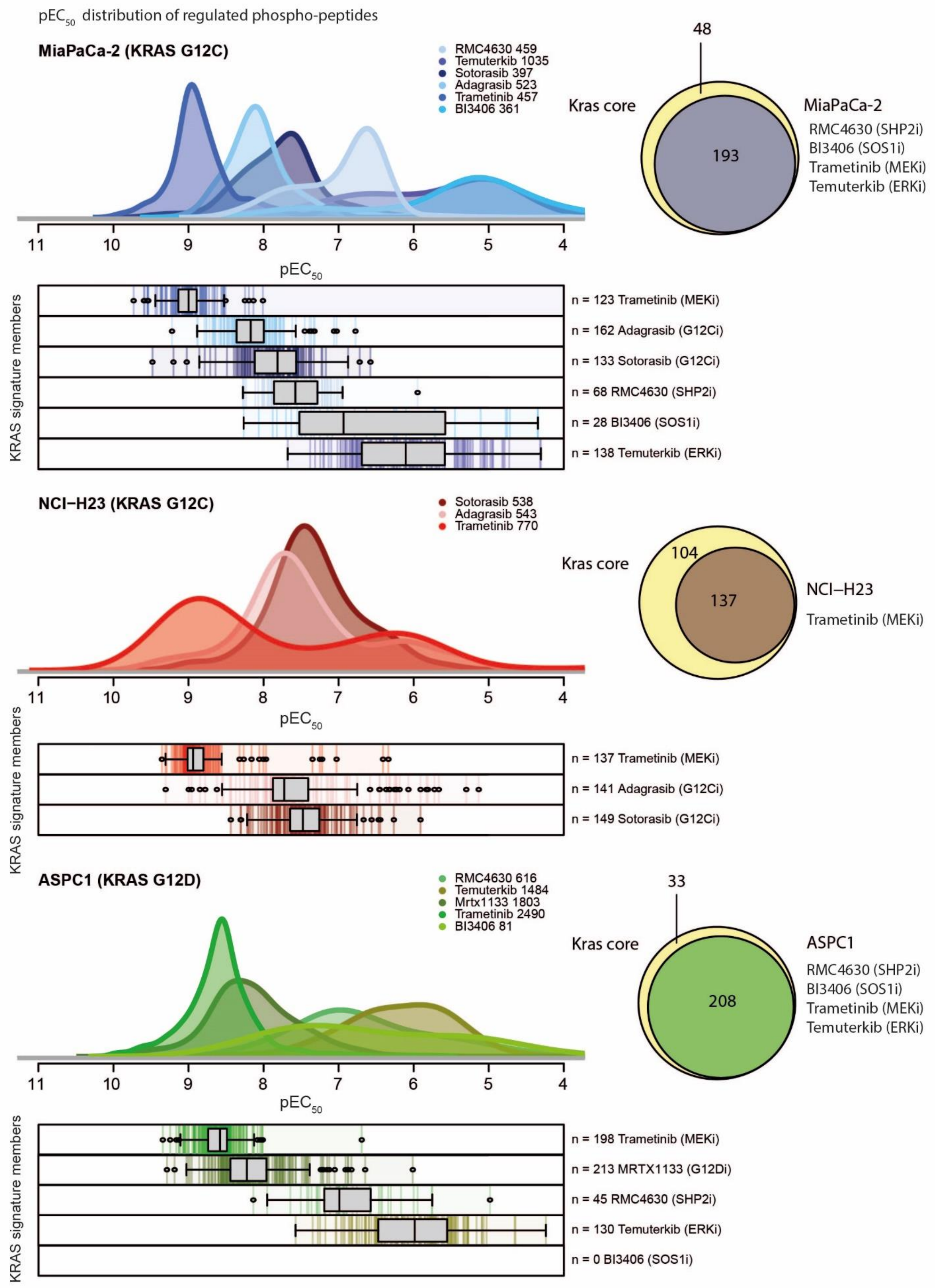

pEC50 distribution of regulated phospho-peptides
MiaPaCa-2 (KRAS G12C)
RMC4630 459
Temuterkib 1035
Sotorasib 397
Adagrasib 523
Trametinib 457
BI3406 361
pEC50
KRAS signature members
n = 123 Trametinib (MEKi)
n = 162 Adagrasib (G12Ci)
n = 133 Sotorasib (G12Ci)
n = 68 RMC4630 (SHP2i)
n = 28 BI3406 (SOS1i)
n = 138 Temuterkib (ERKi)
48
Kras core
193
MiaPaCa-2
RMC4630 (SHP2i)
BI3406 (SOS1i)
Trametinib (MEKi)
Temuterkib (ERKi)
NCI–H23 (KRAS G12C)
Sotorasib 538
Adagrasib 543
Trametinib 770
Kras core
104
137
NCI–H23
Trametinib (MEKi)
pEC50
KRAS signature members
n = 137 Trametinib (MEKi)
n = 141 Adagrasib (G12Ci)
n = 149 Sotorasib (G12Ci)
ASPC1 (KRAS G12D)
RMC4630 616
Temuterkib 1484
Mrtx1133 1803
Trametinib 2490
BI3406 81
33
Kras core
208
ASPC1
RMC4630 (SHP2i)
BI3406 (SOS1i)
Trametinib (MEKi)
Temuterkib (ERKi)
pEC50
KRAS signature members
n = 198 Trametinib (MEKi)
n = 213 MRTX1133 (G12Di)
n = 45 RMC4630 (SHP2i)
n = 130 Temuterkib (ERKi)
n = 0 BI3406 (SOS1i)

**Fig. S11: DecryptM profiling of MAPK pathway inhibitors in mutant KRAS cell lines.**
Distributions of $pEC_{50}$ values of short-term (2h) drug-regulated phospho-peptides for KRASi (Sotorasib, Adagrasib, Mrtx1133), SOSi (BI3406), SHPi (RMC4630), MEKi (Trametinib) and ERKi (Temuterkib) in MiaPaCa-2 (top panel, blue), NCI-H23 (middle panel, red) and ASPC1 (bottom panel, green) cells. Numbers behind drug names indicate the number of phospho-peptides regulated in the experiment. Colored bars in the boxed panels immediately below the distributions show the concentration at which KRAS core signaling signature members were regulated for each drug and cell line. The boxplots summarize the $pEC_{50}$ values from KRAS core signaling signature members (black bar marks the median). Numbers on the right hand side of each boxplot denote the number of phospho-peptides from the KRAS core signaling signature regulated in each data set. Venn diagrams showing how many of the 241 phospho-peptides of the KRAS core signaling signature defined in this study were also drug-regulated by other inhibitors in each of the three cell lines investigated.

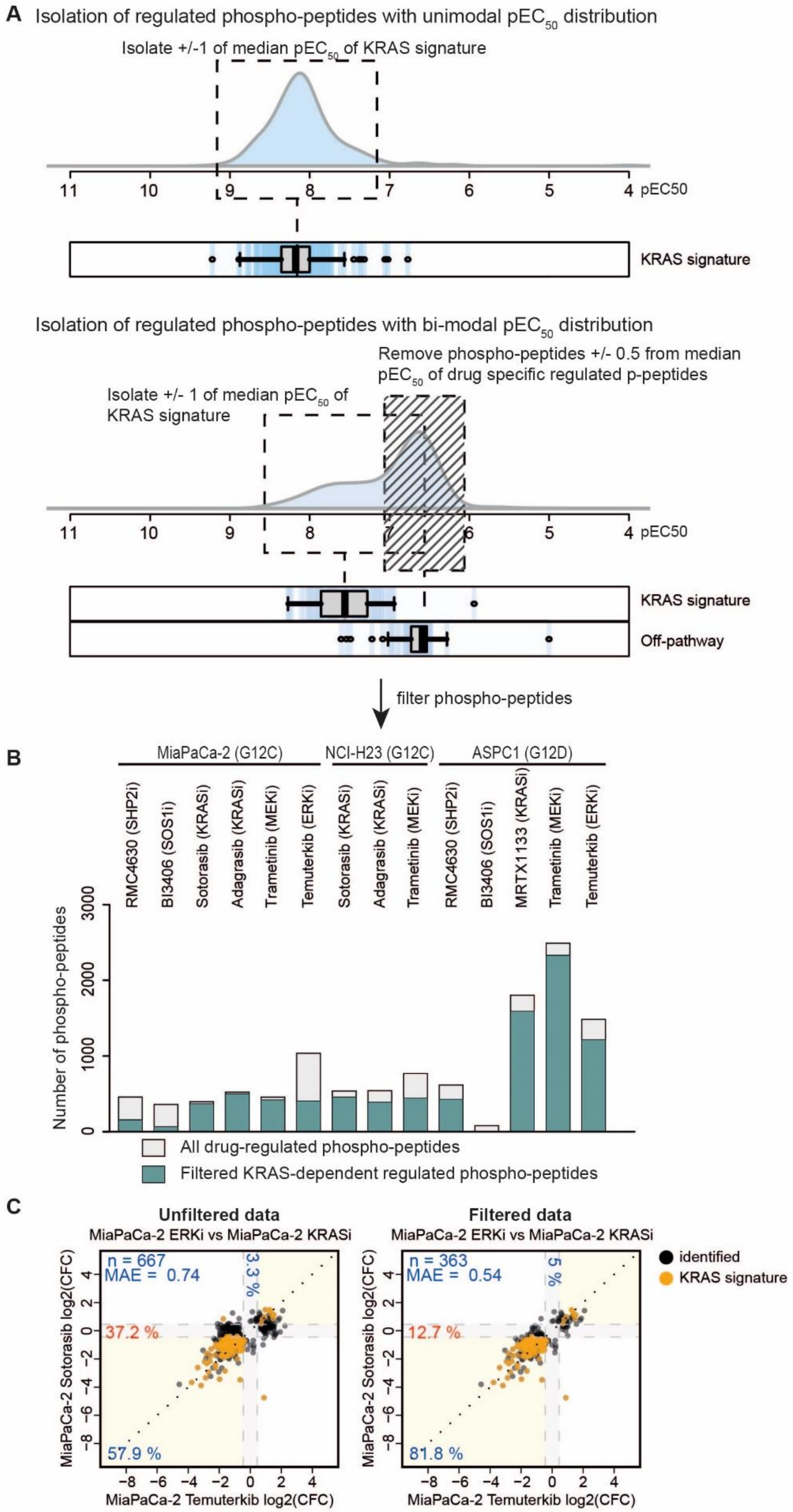
A Isolation of regulated phospho-peptides with unimodal pEC50 distribution
Isolate +/-1 of median pEC50 of KRAS signature
11 10 9 8 7 6 5 4 pEC50
KRAS signature
Isolation of regulated phospho-peptides with bi-modal pEC50 distribution
Remove phospho-peptides +/- 0.5 from median pEC50 of drug specific regulated p-peptides
Isolate +/- 1 of median pEC50 of KRAS signature
11 10 9 8 7 6 5 4 pEC50
KRAS signature
Off-pathway
filter phospho-peptides
B
MiaPaCa-2 (G12C)
NCI-H23 (G12C)
ASPC1 (G12D)
RMC4630 (SHP2i)
BI3406 (SOS1i)
Sotorasib (KRASi)
Adagrasib (KRASi)
Trametinib (MEKi)
Temuterkib (ERKi)
Sotorasib (KRASi)
Adagrasib (KRASi)
Trametinib (MEKi)
RMC4630 (SHP2i)
BI3406 (SOS1i)
MRTX1133 (KRASi)
Trametinib (MEKi)
Temuterkib (ERKi)
Number of phospho-peptides
0 1000 2000 3000
All drug-regulated phospho-peptides
Filtered KRAS-dependent regulated phospho-peptides
C
Unfiltered data
MiaPaCa-2 ERKi vs MiaPaCa-2 KRASi
n = 667
MAE = 0.74
3.3 %
37.2 %
57.9 %
Filtered data
MiaPaCa-2 ERKi vs MiaPaCa-2 KRASi
n = 363
MAE = 0.54
5 %
12.7 %
81.8 %
identified
KRAS signature
MiaPaCa-2 Sotorasib log2(CFC)
MiaPaCa-2 Temuterkib log2(CFC)
−8 −6 −4 −2 0 2 4

**Fig. S12: Separating KRAS pathway members from “off-pathway” effects.**

**(A)** Schematic representation of the approach taken to separate drug-regulated phospho-peptides representing likely KRAS signaling network members from those resulting from “off-target” and “off-pathway” effects. The top distribution illustrates the ideal case that the drug-regulated phospho-peptides show a unimodal distribution of $pEC_{50}$ values based on the assumption that they are all regulated by the same upstream mechanism (i.e. KRAS inhibition). The boxplot below shows the distribution of members of the KRAS core signaling signature defined in this study and regulated within this experiment showing that all but very few fall within ±1 of its median $pEC_{50}$. The panels below show an example of a bimodal $pEC_{50}$ distribution as observed for several drugs in this study and that imply off-target or off-pathway effects. To separate the two underlying distributions, we i) extracted and plotted the $pEC_{50}$ distribution of members of the KRAS core signaling signature and ii) phospho-peptides that were regulated by a single drug within one cell line only (never by any other; termed off-pathway). The median of the latter distribution is close to the apex of the low-potent part of the bimodal distribution, likely containing mostly phospho-peptides that are not part of the KRAS network of phospho-peptides. Only phospho-peptides with $pEC_{50}$ values within ±1 of the median of the KRAS core signaling signature were retained for further analysis. In addition, $pEC_{50}$ values within ±0.5 of the median of the presumed off-target / off-pathway phospho-peptides were removed when bimodal $pEC_{50}$ distributions were observed in the data. We note here that, after filtering, the data may still include a few “drug-specific” phospho-peptides if their $pEC_{50}$ values lie within the area centered around other KRAS signature members. See materials and methods for further details (extended analysis).

**(B)** Barplot showing the number of drug-regulated phospho-peptides before (grey bars) and after (green bars) filtering as described in (A).

**(C)** Scatter plots of pairwise comparisons of ERK inhibition (Temuterkib) vs. KRAS inhibition (Sotorasib) before and after filtering exemplifying the impact of the filtering procedure. The amount of phospho-peptides regulated solely in the MiaPaCa-2 Temuterkib treatment (horizontal grey bar) due to off-target/off-pathway effects was heavily reduced enabling the comparison of the effects of KRAS inhibition vs. ERK inhibition.

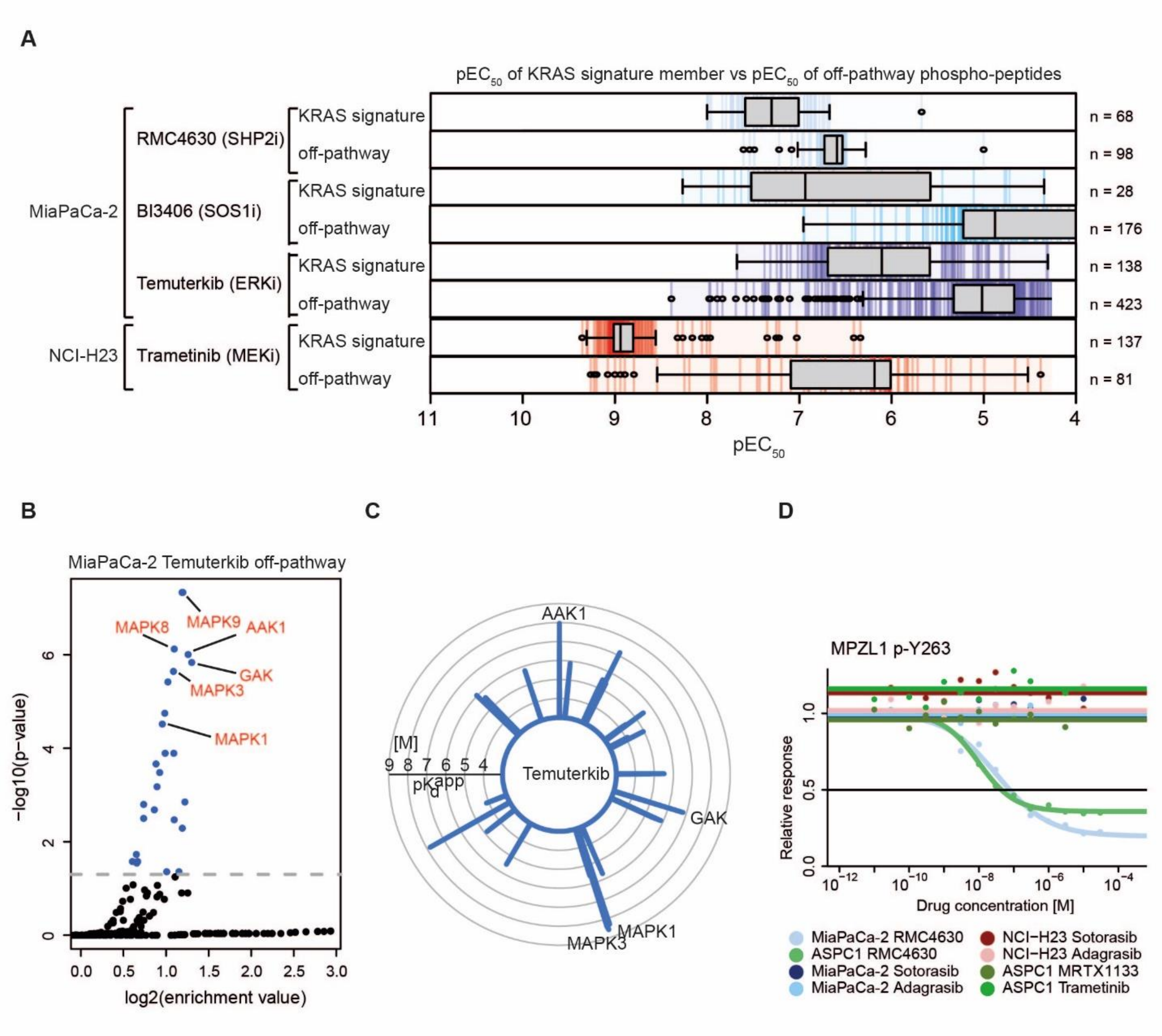


**Fig. S13: Analysis of observed "off-pathway" effects.**

**(A)** Boxplots summarizing the number and potencies of drug-regulated phospho-peptides of KRAS core signaling signature members (upper boxplots) to "off-pathway" phospho-peptides (lower box plots; black bar marks the median $pEC_{50}$) in response to BI3406 (MiaPaCa-2), RMC4630 (MiaPaCa-2), Temuterkib (MiaPaCa-2) and Trametinib (NCI-H23) as described in figure S12.

**(B)** Kinase motif enrichment analysis of Temuterkib off-pathway phospho-peptides. Only phospho-sites with $pEC_{50}$ larger -0.5 from median of off-pathway signature were used for kinase motif enrichment.

**(C)** Radar plot summarizing results of target-deconvolution experiments for Temuterkib using Kinobead pulldowns highlighting several kinase off-targets including some also suggested by the analysis in panel (B; table S10). Every line emerging from the inner circle represents one detected off-target and the length of the line represents the affinity of target engagement (apparent dissociation constant $pK_d^{app}$).

**(D)** Dose-response curves for MPZL1 Y263 phosphorylation in response to SHP2 inhibition. This protein is a putative interaction partner of SHP2 (*63*) and its phosphorylation may, therefore, be used as a proxy for showing target engagement of the SHP2i RMC4630 in MiaPaCa-2 and ASPC1 cells. No effects were observed for KRAS inhibitors, hence, MPZL1 Y263 is not a member of the KRAS core signaling signature.

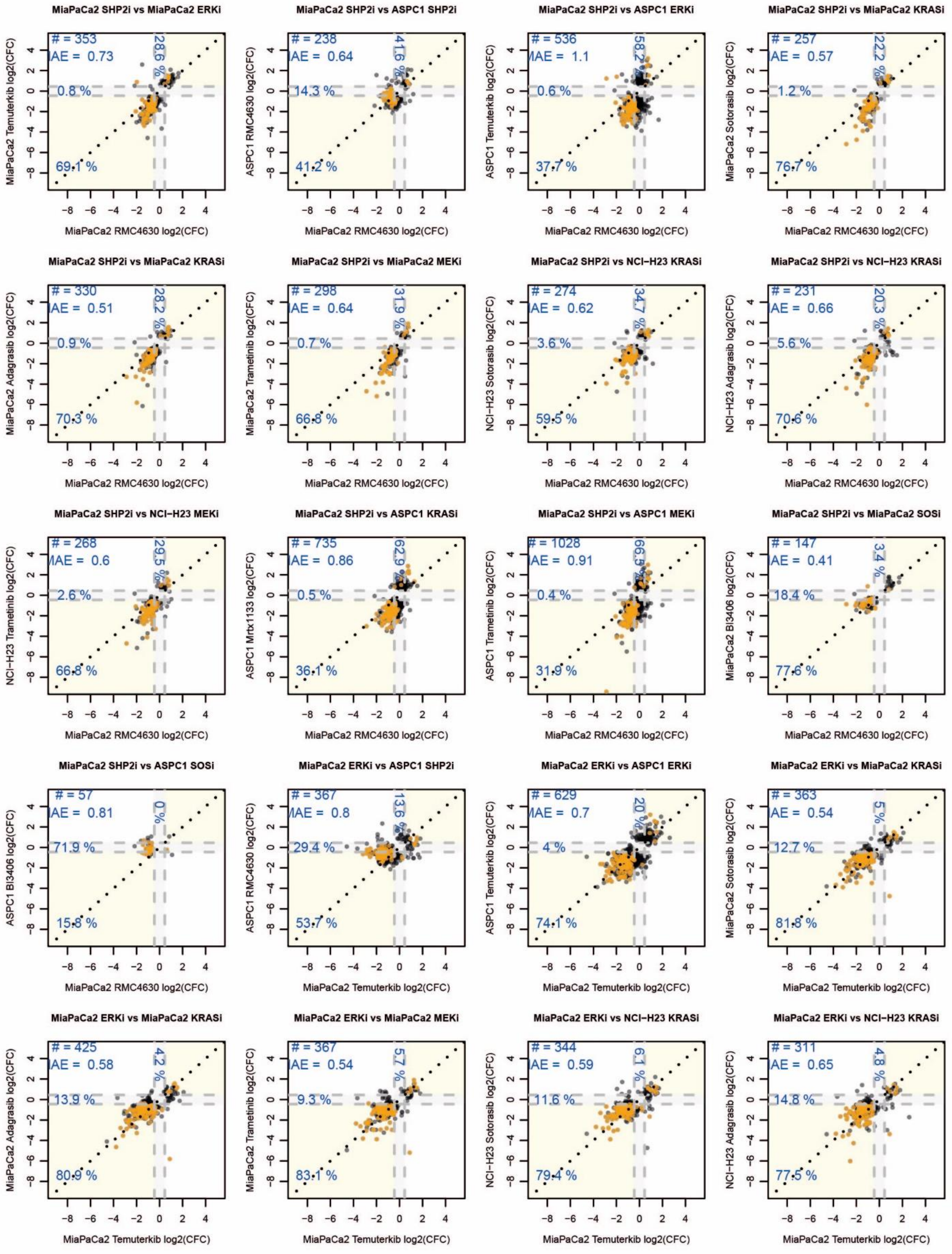
MiaPaCa2 SHP2i vs MiaPaCa2 ERKi
# = 353
AE = 0.73
28.6 %
0.8 %
69.1 %
MiaPaCa2 Temuterkib log2(CFC)
MiaPaCa2 RMC4630 log2(CFC)
MiaPaCa2 SHP2i vs ASPC1 SHP2i
# = 238
AE = 0.64
41.6 %
14.3 %
41.2 %
ASPC1 RMC4630 log2(CFC)
MiaPaCa2 RMC4630 log2(CFC)
MiaPaCa2 SHP2i vs ASPC1 ERKi
# = 536
AE = 1.1
58.2
0.6 %
37.7 %
ASPC1 Temuterkib log2(CFC)
MiaPaCa2 RMC4630 log2(CFC)
MiaPaCa2 SHP2i vs MiaPaCa2 KRASi
# = 257
AE = 0.57
22.2 %
1.2 %
76.7 %
MiaPaCa2 Sotorasib log2(CFC)
MiaPaCa2 RMC4630 log2(CFC)
MiaPaCa2 SHP2i vs MiaPaCa2 KRASi
# = 330
AE = 0.51
28.2 %
0.9 %
70.3 %
MiaPaCa2 Adagrasib log2(CFC)
MiaPaCa2 RMC4630 log2(CFC)
MiaPaCa2 SHP2i vs MiaPaCa2 MEKi
# = 298
AE = 0.64
31.9 %
0.7 %
66.8 %
MiaPaCa2 Trametinib log2(CFC)
MiaPaCa2 RMC4630 log2(CFC)
MiaPaCa2 SHP2i vs NCI−H23 KRASi
# = 274
AE = 0.62
34.7 %
3.6 %
59.5 %
NCI−H23 Sotorasib log2(CFC)
MiaPaCa2 RMC4630 log2(CFC)
MiaPaCa2 SHP2i vs NCI−H23 KRASi
# = 231
AE = 0.66
20.3 %
5.6 %
70.6 %
NCI−H23 Adagrasib log2(CFC)
MiaPaCa2 RMC4630 log2(CFC)
MiaPaCa2 SHP2i vs NCI−H23 MEKi
# = 268
MAE = 0.6
29.5 %
2.6 %
66.8 %
NCI−H23 Trametinib log2(CFC)
MiaPaCa2 RMC4630 log2(CFC)
MiaPaCa2 SHP2i vs ASPC1 KRASi
# = 735
AE = 0.86
62.9
0.5 %
36.1 %
ASPC1 Mrtx1133 log2(CFC)
MiaPaCa2 RMC4630 log2(CFC)
MiaPaCa2 SHP2i vs ASPC1 MEKi
# = 1028
MAE = 0.91
66.5
0.4 %
31.9 %
ASPC1 Trametinib log2(CFC)
MiaPaCa2 RMC4630 log2(CFC)
MiaPaCa2 SHP2i vs MiaPaCa2 SOSi
# = 147
AE = 0.41
3.4 %
18.4 %
77.6 %
MiaPaCa2 BI3406 log2(CFC)
MiaPaCa2 RMC4630 log2(CFC)
MiaPaCa2 SHP2i vs ASPC1 SOSi
# = 57
AE = 0.81
0 %
71.9 %
15.8 %
ASPC1 BI3406 log2(CFC)
MiaPaCa2 RMC4630 log2(CFC)
MiaPaCa2 ERKi vs ASPC1 SHP2i
# = 367
MAE = 0.8
13.6
29.4 %
53.7 %
ASPC1 RMC4630 log2(CFC)
MiaPaCa2 Temuterkib log2(CFC)
MiaPaCa2 ERKi vs ASPC1 ERKi
# = 629
MAE = 0.7
20 %
4 %
74.1 %
ASPC1 Temuterkib log2(CFC)
MiaPaCa2 Temuterkib log2(CFC)
MiaPaCa2 ERKi vs MiaPaCa2 KRASi
# = 363
AE = 0.54
5 %
12.7 %
81.8 %
MiaPaCa2 Sotorasib log2(CFC)
MiaPaCa2 Temuterkib log2(CFC)
MiaPaCa2 ERKi vs MiaPaCa2 KRASi
# = 425
AE = 0.58
4.2 %
13.9 %
80.9 %
MiaPaCa2 Adagrasib log2(CFC)
MiaPaCa2 Temuterkib log2(CFC)
MiaPaCa2 ERKi vs MiaPaCa2 MEKi
# = 367
AE = 0.54
5.7 %
9.3 %
83.1 %
MiaPaCa2 Trametinib log2(CFC)
MiaPaCa2 Temuterkib log2(CFC)
MiaPaCa2 ERKi vs NCI−H23 KRASi
# = 344
AE = 0.59
6.1 %
11.6 %
79.4 %
NCI−H23 Sotorasib log2(CFC)
MiaPaCa2 Temuterkib log2(CFC)
MiaPaCa2 ERKi vs NCI−H23 KRASi
# = 311
AE = 0.65
4.8 %
14.8 %
77.5 %
NCI−H23 Adagrasib log2(CFC)
MiaPaCa2 Temuterkib log2(CFC)

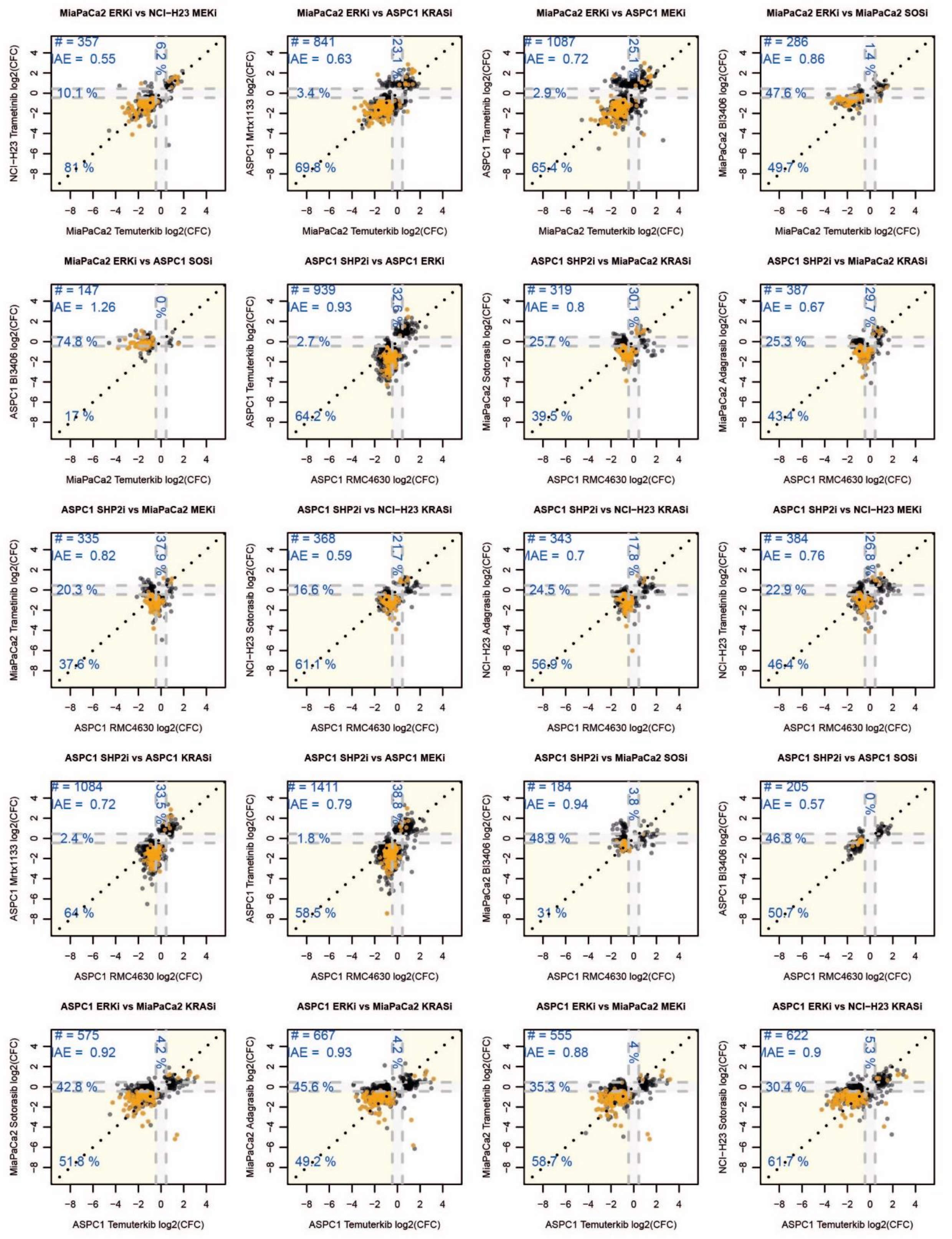
MiaPaCa2 ERKi vs NCI−H23 MEKi
# = 357
MAE = 0.55
6.2 %
10.1 %
81 %
NCI−H23 Trametinib log2(CFC)
MiaPaCa2 Temuterkib log2(CFC)
MiaPaCa2 ERKi vs ASPC1 KRASi
# = 841
MAE = 0.63
23.1 %
3.4 %
69.8 %
ASPC1 Mrtx1133 log2(CFC)
MiaPaCa2 Temuterkib log2(CFC)
MiaPaCa2 ERKi vs ASPC1 MEKi
# = 1087
MAE = 0.72
25.1 %
2.9 %
65.4 %
ASPC1 Trametinib log2(CFC)
MiaPaCa2 Temuterkib log2(CFC)
MiaPaCa2 ERKi vs MiaPaCa2 SOSi
# = 286
MAE = 0.86
1.4 %
47.6 %
49.7 %
MiaPaCa2 BI3406 log2(CFC)
MiaPaCa2 Temuterkib log2(CFC)
MiaPaCa2 ERKi vs ASPC1 SOSi
# = 147
MAE = 1.26
0 %
74.8 %
17 %
ASPC1 BI3406 log2(CFC)
MiaPaCa2 Temuterkib log2(CFC)
ASPC1 SHP2i vs ASPC1 ERKi
# = 939
MAE = 0.93
32.6 %
2.7 %
64.2 %
ASPC1 Temuterkib log2(CFC)
ASPC1 RMC4630 log2(CFC)
ASPC1 SHP2i vs MiaPaCa2 KRASi
# = 319
MAE = 0.8
30.1 %
25.7 %
39.5 %
MiaPaCa2 Sotorasib log2(CFC)
ASPC1 RMC4630 log2(CFC)
ASPC1 SHP2i vs MiaPaCa2 KRASi
# = 387
MAE = 0.67
29.7 %
25.3 %
43.4 %
MiaPaCa2 Adagrasib log2(CFC)
ASPC1 RMC4630 log2(CFC)
ASPC1 SHP2i vs MiaPaCa2 MEKi
# = 335
MAE = 0.82
37.9 %
20.3 %
37.6 %
MiaPaCa2 Trametinib log2(CFC)
ASPC1 RMC4630 log2(CFC)
ASPC1 SHP2i vs NCI−H23 KRASi
# = 368
MAE = 0.59
21.7 %
16.6 %
61.1 %
NCI−H23 Sotorasib log2(CFC)
ASPC1 RMC4630 log2(CFC)
ASPC1 SHP2i vs NCI−H23 KRASi
# = 343
MAE = 0.7
17.8 %
24.5 %
56.9 %
NCI−H23 Adagrasib log2(CFC)
ASPC1 RMC4630 log2(CFC)
ASPC1 SHP2i vs NCI−H23 MEKi
# = 384
MAE = 0.76
26.8 %
22.9 %
46.4 %
NCI−H23 Trametinib log2(CFC)
ASPC1 RMC4630 log2(CFC)
ASPC1 SHP2i vs ASPC1 KRASi
# = 1084
MAE = 0.72
33.5 %
2.4 %
64 %
ASPC1 Mrtx1133 log2(CFC)
ASPC1 RMC4630 log2(CFC)
ASPC1 SHP2i vs ASPC1 MEKi
# = 1411
MAE = 0.79
38.8 %
1.8 %
58.5 %
ASPC1 Trametinib log2(CFC)
ASPC1 RMC4630 log2(CFC)
ASPC1 SHP2i vs MiaPaCa2 SOSi
# = 184
MAE = 0.94
3.8 %
48.9 %
31 %
MiaPaCa2 BI3406 log2(CFC)
ASPC1 RMC4630 log2(CFC)
ASPC1 SHP2i vs ASPC1 SOSi
# = 205
MAE = 0.57
0 %
46.8 %
50.7 %
ASPC1 BI3406 log2(CFC)
ASPC1 RMC4630 log2(CFC)
ASPC1 ERKi vs MiaPaCa2 KRASi
# = 575
MAE = 0.92
4.2 %
42.8 %
51.8 %
MiaPaCa2 Sotorasib log2(CFC)
ASPC1 Temuterkib log2(CFC)
ASPC1 ERKi vs MiaPaCa2 KRASi
# = 667
MAE = 0.93
4.2 %
45.6 %
49.2 %
MiaPaCa2 Adagrasib log2(CFC)
ASPC1 Temuterkib log2(CFC)
ASPC1 ERKi vs MiaPaCa2 MEKi
# = 555
MAE = 0.88
4 %
35.3 %
58.7 %
MiaPaCa2 Trametinib log2(CFC)
ASPC1 Temuterkib log2(CFC)
ASPC1 ERKi vs NCI−H23 KRASi
# = 622
MAE = 0.9
5.3 %
30.4 %
61.7 %
NCI−H23 Sotorasib log2(CFC)
ASPC1 Temuterkib log2(CFC)

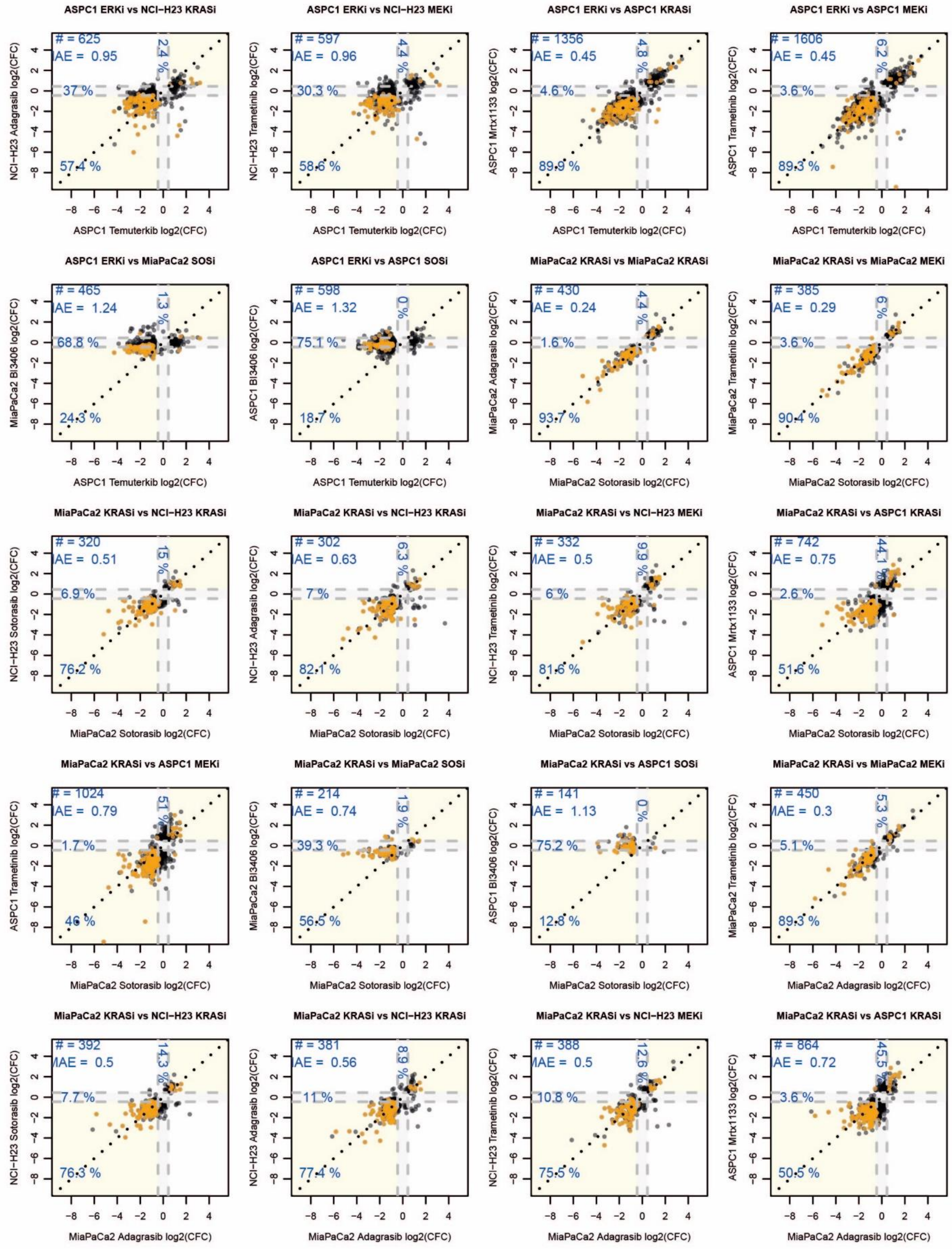
ASPC1 ERKi vs NCI−H23 KRASi
# = 625
AE = 0.95
2.4 %
37 %
57.4 %
NCI−H23 Adagrasib log2(CFC)
ASPC1 Temuterkib log2(CFC)
ASPC1 ERKi vs NCI−H23 MEKi
# = 597
AE = 0.96
4.4 %
30.3 %
58.6 %
NCI−H23 Trametinib log2(CFC)
ASPC1 Temuterkib log2(CFC)
ASPC1 ERKi vs ASPC1 KRASi
# = 1356
AE = 0.45
4.8 %
4.6 %
89.9 %
ASPC1 Mrtx1133 log2(CFC)
ASPC1 Temuterkib log2(CFC)
ASPC1 ERKi vs ASPC1 MEKi
# = 1606
AE = 0.45
6.2 %
3.6 %
89.3 %
ASPC1 Trametinib log2(CFC)
ASPC1 Temuterkib log2(CFC)
ASPC1 ERKi vs MiaPaCa2 SOSi
# = 465
AE = 1.24
1.3 %
68.8 %
24.3 %
MiaPaCa2 BI3406 log2(CFC)
ASPC1 Temuterkib log2(CFC)
ASPC1 ERKi vs ASPC1 SOSi
# = 598
AE = 1.32
0 %
75.1 %
18.7 %
ASPC1 BI3406 log2(CFC)
ASPC1 Temuterkib log2(CFC)
MiaPaCa2 KRASi vs MiaPaCa2 KRASi
# = 430
AE = 0.24
4.4 %
1.6 %
93.7 %
MiaPaCa2 Adagrasib log2(CFC)
MiaPaCa2 Sotorasib log2(CFC)
MiaPaCa2 KRASi vs MiaPaCa2 MEKi
# = 385
AE = 0.29
6 %
3.6 %
90.4 %
MiaPaCa2 Trametinib log2(CFC)
MiaPaCa2 Sotorasib log2(CFC)
MiaPaCa2 KRASi vs NCI−H23 KRASi
# = 320
AE = 0.51
15 %
6.9 %
76.2 %
NCI−H23 Sotorasib log2(CFC)
MiaPaCa2 Sotorasib log2(CFC)
MiaPaCa2 KRASi vs NCI−H23 KRASi
# = 302
AE = 0.63
6.3 %
7 %
82.1 %
NCI−H23 Adagrasib log2(CFC)
MiaPaCa2 Sotorasib log2(CFC)
MiaPaCa2 KRASi vs NCI−H23 MEKi
# = 332
MAE = 0.5
9.9 %
6 %
81.6 %
NCI−H23 Trametinib log2(CFC)
MiaPaCa2 Sotorasib log2(CFC)
MiaPaCa2 KRASi vs ASPC1 KRASi
# = 742
AE = 0.75
44.1 %
2.6 %
51.6 %
ASPC1 Mrtx1133 log2(CFC)
MiaPaCa2 Sotorasib log2(CFC)
MiaPaCa2 KRASi vs ASPC1 MEKi
# = 1024
AE = 0.79
51 %
1.7 %
46 %
ASPC1 Trametinib log2(CFC)
MiaPaCa2 Sotorasib log2(CFC)
MiaPaCa2 KRASi vs MiaPaCa2 SOSi
# = 214
AE = 0.74
1.9 %
39.3 %
56.5 %
MiaPaCa2 BI3406 log2(CFC)
MiaPaCa2 Sotorasib log2(CFC)
MiaPaCa2 KRASi vs ASPC1 SOSi
# = 141
AE = 1.13
0 %
75.2 %
12.8 %
ASPC1 BI3406 log2(CFC)
MiaPaCa2 Sotorasib log2(CFC)
MiaPaCa2 KRASi vs MiaPaCa2 MEKi
# = 450
MAE = 0.3
5.3 %
5.1 %
89.3 %
MiaPaCa2 Trametinib log2(CFC)
MiaPaCa2 Adagrasib log2(CFC)
MiaPaCa2 KRASi vs NCI−H23 KRASi
# = 392
MAE = 0.5
14.3 %
7.7 %
76.3 %
NCI−H23 Sotorasib log2(CFC)
MiaPaCa2 Adagrasib log2(CFC)
MiaPaCa2 KRASi vs NCI−H23 KRASi
# = 381
AE = 0.56
8.9 %
11 %
77.4 %
NCI−H23 Adagrasib log2(CFC)
MiaPaCa2 Adagrasib log2(CFC)
MiaPaCa2 KRASi vs NCI−H23 MEKi
# = 388
MAE = 0.5
12.6 %
10.8 %
75.5 %
NCI−H23 Trametinib log2(CFC)
MiaPaCa2 Adagrasib log2(CFC)
MiaPaCa2 KRASi vs ASPC1 KRASi
# = 864
AE = 0.72
45.5 %
3.6 %
50.5 %
ASPC1 Mrtx1133 log2(CFC)
MiaPaCa2 Adagrasib log2(CFC)

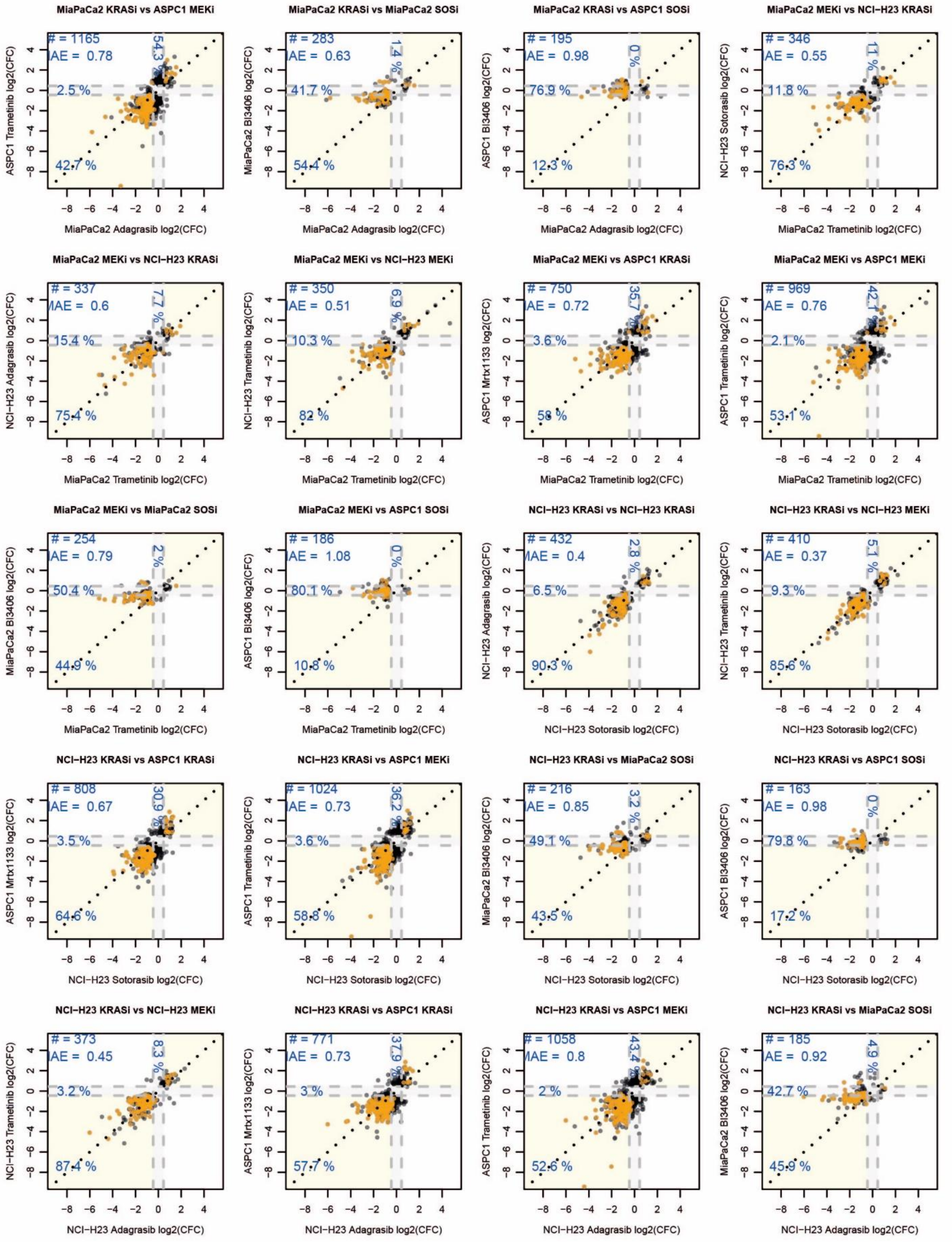

MiaPaCa2 KRASi vs ASPC1 MEKi
# = 1165
AE = 0.78
54.3 %
2.5 %
42.7 %
ASPC1 Trametinib log2(CFC)
MiaPaCa2 Adagrasib log2(CFC)
MiaPaCa2 KRASi vs MiaPaCa2 SOSi
# = 283
AE = 0.63
1.4 %
41.7 %
54.4 %
MiaPaCa2 BI3406 log2(CFC)
MiaPaCa2 Adagrasib log2(CFC)
MiaPaCa2 KRASi vs ASPC1 SOSi
# = 195
AE = 0.98
0 %
76.9 %
12.3 %
ASPC1 BI3406 log2(CFC)
MiaPaCa2 Adagrasib log2(CFC)
MiaPaCa2 MEKi vs NCI−H23 KRASi
# = 346
AE = 0.55
11 %
11.8 %
76.3 %
NCI−H23 Sotorasib log2(CFC)
MiaPaCa2 Trametinib log2(CFC)
MiaPaCa2 MEKi vs NCI−H23 KRASi
# = 337
AE = 0.6
7.7 %
15.4 %
75.4 %
NCI−H23 Adagrasib log2(CFC)
MiaPaCa2 Trametinib log2(CFC)
MiaPaCa2 MEKi vs NCI−H23 MEKi
# = 350
AE = 0.51
6.9 %
10.3 %
82 %
NCI−H23 Trametinib log2(CFC)
MiaPaCa2 Trametinib log2(CFC)
MiaPaCa2 MEKi vs ASPC1 KRASi
# = 750
AE = 0.72
35.7 %
3.6 %
58 %
ASPC1 Mrtx1133 log2(CFC)
MiaPaCa2 Trametinib log2(CFC)
MiaPaCa2 MEKi vs ASPC1 MEKi
# = 969
AE = 0.76
42.1 %
2.1 %
53.1 %
ASPC1 Trametinib log2(CFC)
MiaPaCa2 Trametinib log2(CFC)
MiaPaCa2 MEKi vs MiaPaCa2 SOSi
# = 254
AE = 0.79
2 %
50.4 %
44.9 %
MiaPaCa2 BI3406 log2(CFC)
MiaPaCa2 Trametinib log2(CFC)
MiaPaCa2 MEKi vs ASPC1 SOSi
# = 186
AE = 1.08
0 %
80.1 %
10.8 %
ASPC1 BI3406 log2(CFC)
MiaPaCa2 Trametinib log2(CFC)
NCI−H23 KRASi vs NCI−H23 KRASi
# = 432
AE = 0.4
2.8 %
6.5 %
90.3 %
NCI−H23 Adagrasib log2(CFC)
NCI−H23 Sotorasib log2(CFC)
NCI−H23 KRASi vs NCI−H23 MEKi
# = 410
AE = 0.37
5.1 %
9.3 %
85.6 %
NCI−H23 Trametinib log2(CFC)
NCI−H23 Sotorasib log2(CFC)
NCI−H23 KRASi vs ASPC1 KRASi
# = 808
AE = 0.67
30.9 %
3.5 %
64.6 %
ASPC1 Mrtx1133 log2(CFC)
NCI−H23 Sotorasib log2(CFC)
NCI−H23 KRASi vs ASPC1 MEKi
# = 1024
AE = 0.73
36.2 %
3.6 %
58.8 %
ASPC1 Trametinib log2(CFC)
NCI−H23 Sotorasib log2(CFC)
NCI−H23 KRASi vs MiaPaCa2 SOSi
# = 216
AE = 0.85
3.2 %
49.1 %
43.5 %
MiaPaCa2 BI3406 log2(CFC)
NCI−H23 Sotorasib log2(CFC)
NCI−H23 KRASi vs ASPC1 SOSi
# = 163
AE = 0.98
0 %
79.8 %
17.2 %
ASPC1 BI3406 log2(CFC)
NCI−H23 Sotorasib log2(CFC)
NCI−H23 KRASi vs NCI−H23 MEKi
# = 373
AE = 0.45
8.3 %
3.2 %
87.4 %
NCI−H23 Trametinib log2(CFC)
NCI−H23 Adagrasib log2(CFC)
NCI−H23 KRASi vs ASPC1 KRASi
# = 771
AE = 0.73
37.9 %
3 %
57.7 %
ASPC1 Mrtx1133 log2(CFC)
NCI−H23 Adagrasib log2(CFC)
NCI−H23 KRASi vs ASPC1 MEKi
# = 1058
AE = 0.8
43.4 %
2 %
52.6 %
ASPC1 Trametinib log2(CFC)
NCI−H23 Adagrasib log2(CFC)
NCI−H23 KRASi vs MiaPaCa2 SOSi
# = 185
AE = 0.92
4.9 %
42.7 %
45.9 %
MiaPaCa2 BI3406 log2(CFC)
NCI−H23 Adagrasib log2(CFC)

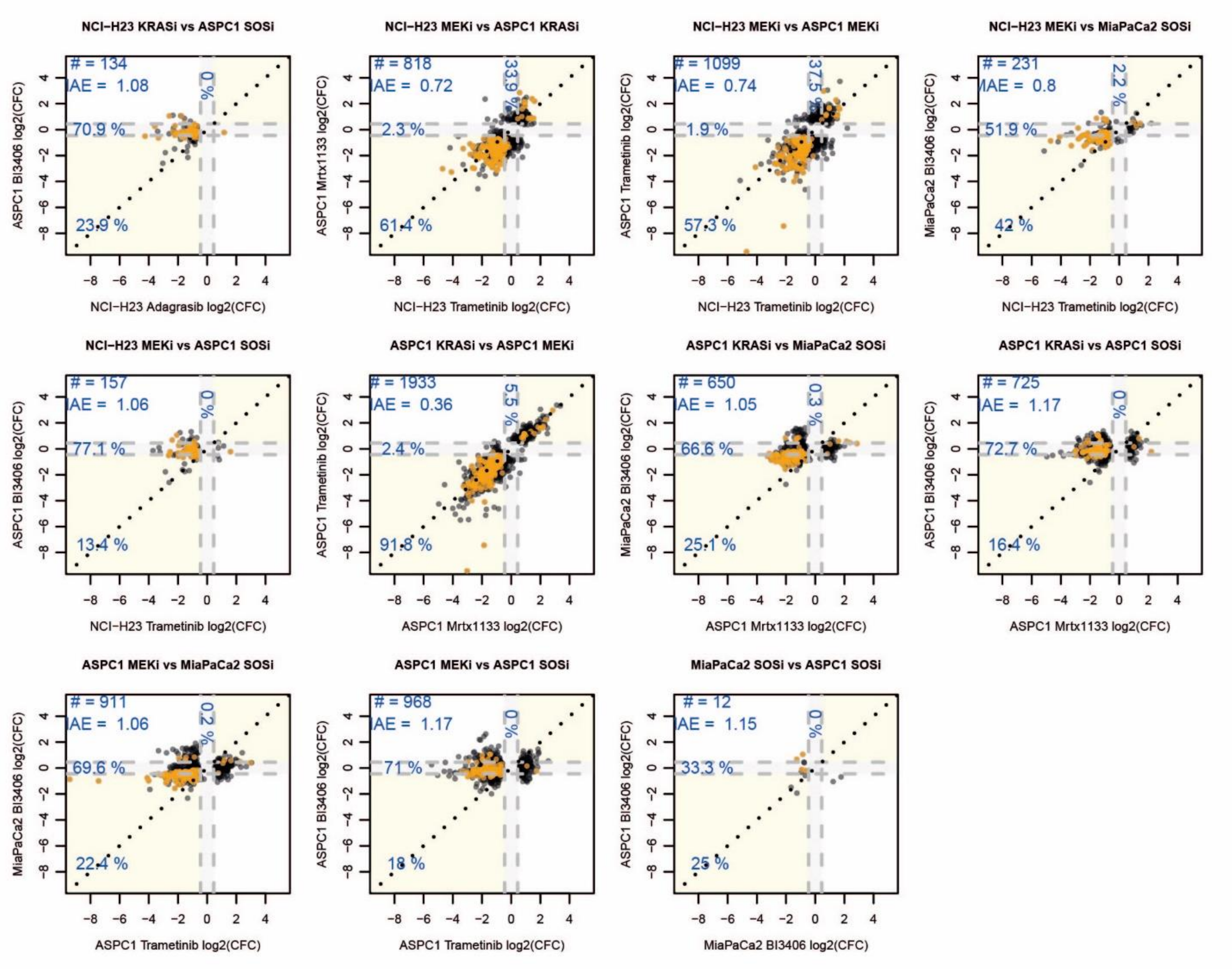


**Fig. S14: Comparison of decryptM profiles from different MAPK-pathway inhibiting drugs.** Scatter plots showing the pairwise comparison of drug-regulated phospho-peptide responses for any combination of KRAS inhibitor and cell line from the heatmap shown in main manuscript Fig. 5C. MAE: median absolute error calculated from log2 curve fold changes; # number of phospho-peptides in plot; Dashed lines mark the boundaries of CurveCurator log2 fold change cut-off (fc-values = ±0.45). Yellow areas contain phospho-peptides with consistent responses, grey areas contain phospho-peptides regulated by one of the conditions only; Percentages denote the relative proportion of phospho-peptides in the yellow and grey areas respectively. Values on the y- and x-axis are the log2 curve fold change (CFC) of phospho-peptides in response to drug; Orange points mark phospho-peptides from the KRAS core signature.

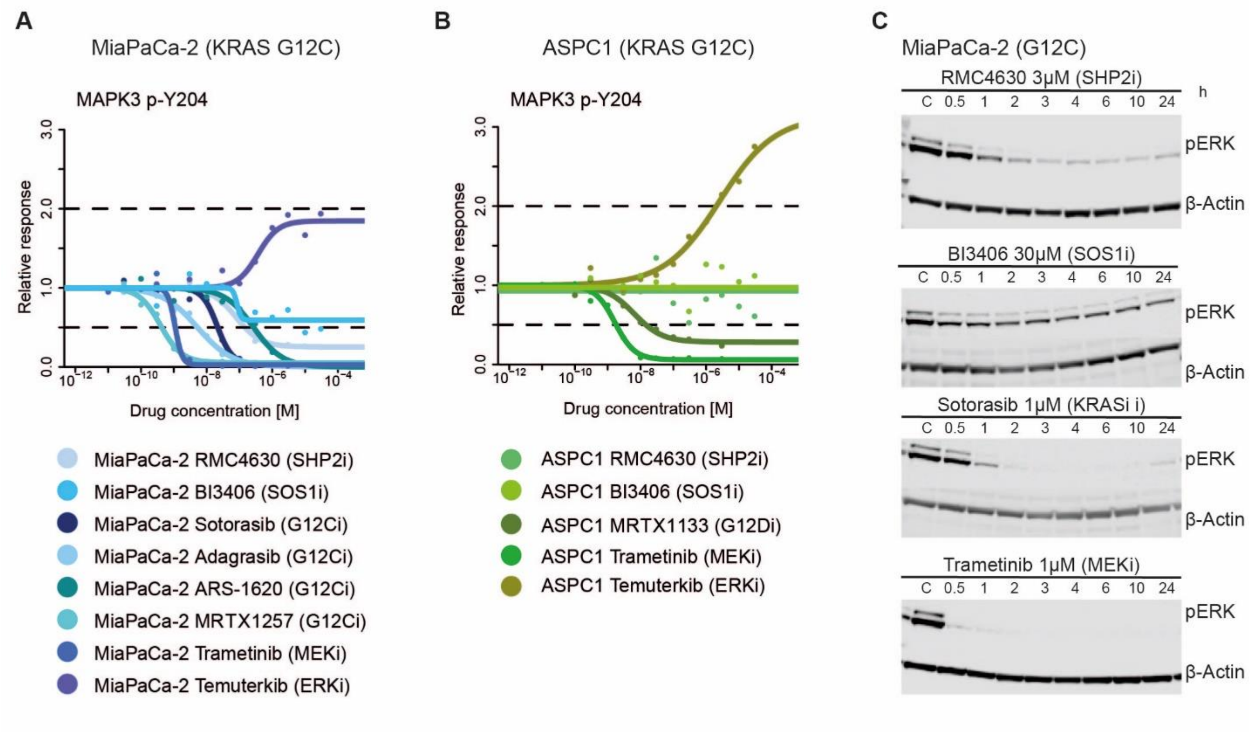


**Fig. S15: Effects of different MAPK-pathway modulating drugs on ERK activation state**
**(A)** Dose-response curves of MAPK3 Y204 (activation loop of MAPK3) phosphorylation in response to drugs acting upstream or downstream of KRAS (2h; MiaPaCa-2 cells). We note that the apparent activation of MAPK3 (p-Y204) upon Temuterkib is actually just accumulation of phosphorylation as the result of upstream MEK activity (*64*). Because MAPK3 is inhibited by Temuterkib, all downstream signaling is abrogated.
**(B)** Same as (A) but for ASPC1 cells.
**(C)** Western blot analysis of phospho-MAPK1/3 in response to SHP2, SOS1, KRAS and MEK inhibition over the course of 24h (C = control). We note that SOS1 and SHP2 inhibitors do not fully abrogate MAPK activity while KRAS and MEK inhibition does.

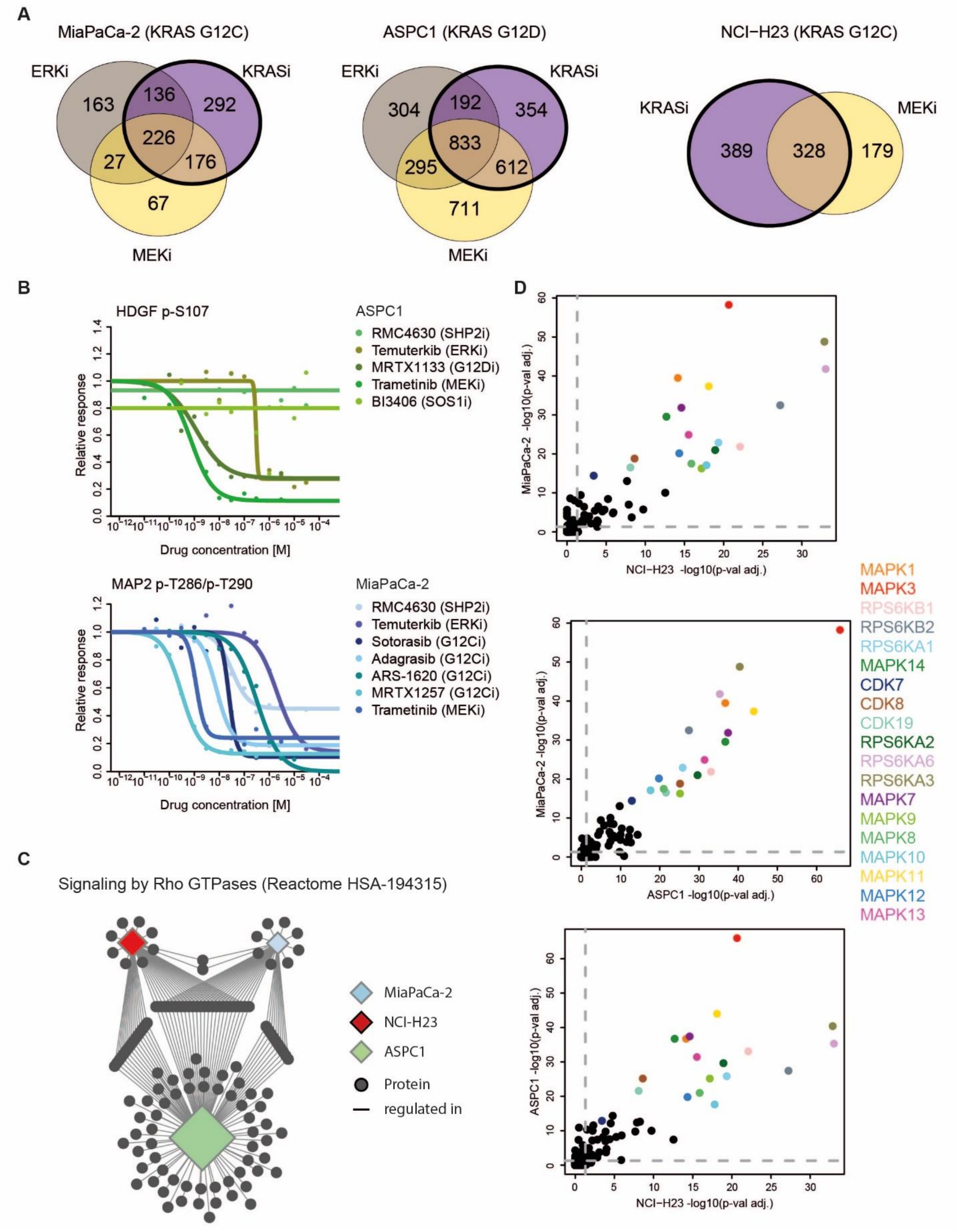


**Fig. S16: Cell line-specific KRAS network.**

**(A)** Venn diagrams comparing the number of drug-regulated phospho-peptides in a particular cell line in response to KRAS inhibitors and downstream MAPK pathway-modulating drugs (ERK and MEK inhibition). Phospho-peptides were curated per cell line. Regulated phospho-peptides were included in the cell-specific KRAS signature network if they were meeting filter criteria (fig. S12) in all three inhibitor classes (KRASi, MEKi, ERKi) within one cell line. Regulated phospho-peptides that met filter criteria (fig. S12) in at least one but not all drug treatments within one cell line (because the required fold change response was not quite reached) were included in the KRAS network within this cell line, as long as they showed a CurveCurator p-value of $\leq 0.01$, an absolute log2 fold change (response) of $\geq 0.6$ and followed the same quality criteria as regular curves ($EC_{50}$ must lie within the treatment range, see materials and methods (classification) for details).
**(B)** Example dose-response curves of phospho-peptides that showed consistent drug regulation in one cell line, but never in another.
**(C)** Network of proteins representing drug-regulated phospho-peptides in response to KRAS inhibition and that are annotated members of Rho GTPase signaling (HSA-194315, Reactome). Diamond node sizes represents the relative statistical significance (adjusted p-val.) of the enriched pathway in the respective cell line.
**(D)** Pair-wise comparison of enriched Kinase motifs between cell lines from analysis of down-regulated phospho-peptides. We note that despite substantial differences in number of drug-regulated phospho-peptides detected in each cell line, the same kinase motifs dominate the response to inhibition of the KRAS-MEK-MAPK axis (KRASi data only).

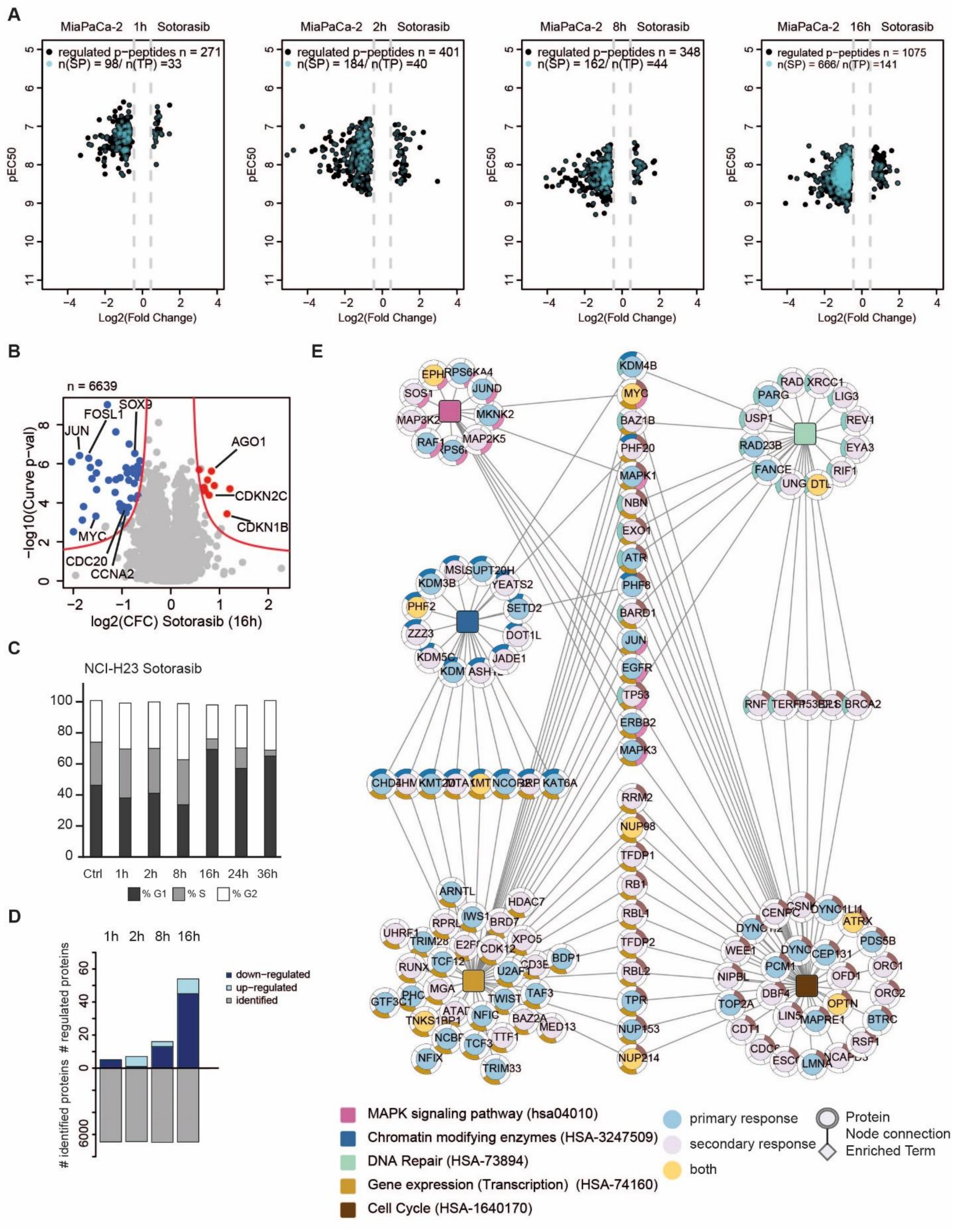
A
MiaPaCa-2 1h Sotorasib
regulated p-peptides n = 271
n(SP) = 98/ n(TP) =33
MiaPaCa-2 2h Sotorasib
regulated p-peptides n = 401
n(SP) = 184/ n(TP) =40
MiaPaCa-2 8h Sotorasib
regulated p-peptides n = 348
n(SP) = 162/ n(TP) =44
MiaPaCa-2 16h Sotorasib
regulated p-peptides n = 1075
n(SP) = 666/ n(TP) =141
pEC50
Log2(Fold Change)
B
n = 6639
JUN
FOSL1
SOX9
AGO1
CDKN2C
CDKN1B
MYC
CDC20
CCNA2
-log10(Curve p-val)
log2(CFC) Sotorasib (16h)
C
NCI-H23 Sotorasib
Ctrl 1h 2h 8h 16h 24h 36h
% G1 % S % G2
D
1h 2h 8h 16h
down-regulated
up-regulated
identified
### regulated proteins
### identified proteins
E
MAPK signaling pathway (hsa04010)
Chromatin modifying enzymes (HSA-3247509)
DNA Repair (HSA-73894)
Gene expression (Transcription) (HSA-74160)
Cell Cycle (HSA-1640170)
primary response
secondary response
both
Protein
Node connection
Enriched Term

**Fig. S17: 2D decryptM analysis of Sotorasib in MiaPaCa-2 cells**

**(A)** Scatter plots showing the potencies ($pEC_{50}$) and log2 responses (log2 curve fold change) of regulated phospho-peptides upon Sotorasib treatment of MiaPaCa-2 cells after 1, 2, 8 and 16h (filtered for phospho-peptides with $pEC_{50}$ ± 1 of median $pEC_{50}$ KRAS signature members; fig. S12). SP/TP motif containing phospho-peptides are highlighted in cyan and their proportion increases over time.

**(B)** Volcano plot showing results of dose-dependent protein expression changes after 16h of Sotorasib treatment in MiaPaCa-2 cells. Down-regulated protein groups (CurveCurator criteria plus outlier filter; see materials and methods (classification) for details) are highlighted in blue and up-regulated in red. Red asymptote marks the boundary for curve classification based on CurveCurator.

**(C)** Barplot showing the proportion of NCI-H23 cells in different stages of the cell cycle determined by FACS analysis of propidium iodide (PI) stained cells following Sotorasib treatment (0-36h, control = Ctrl).

**(D)** Barplot showing the number of identified proteins groups and drug-regulated protein abundance changes of Sotorasib-treated MiaPaCa-2 cells.

**(E)** Functional enrichment of protein coding genes (representing the underlying drug-regulated phospho-peptides) by membership in different functional categories. Proteins in blue represent phospho-peptides that are part of the immediate response, in pink those of the adaptive response and in orange, those that are part of both.

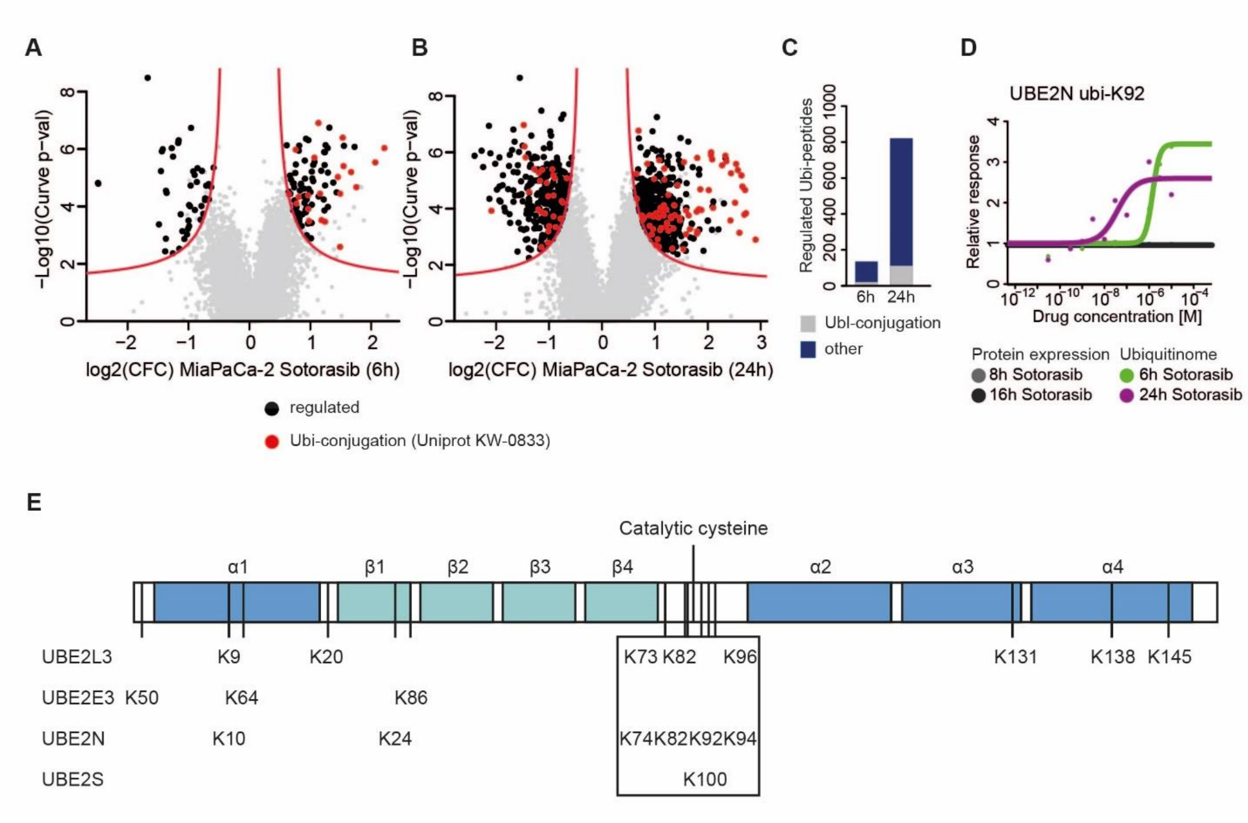


**Fig. S18: Effects of KRAS inhibition on the ubiquitinome**

**(A, B)** Volcano plots of ubiquitinylated peptides after 6h (A) and 24h (B) of Sotorasib-treated MiaPaCa-2 cells. Drug-regulated ubiquitinylated peptides are highlighted in black and drug-regulated ubiquitinylated peptides associated with Ubl-conjugation (according to UniProt(KW-0833) in red.

**(C)** Barplot summarizing the data shown in panels (A) and (B).

**(D)** Dose-response curve of ubiquitylation of UBE2N K92 and protein expression of UBE2N.

**(E)** Schematic representation of E2 ligase domain structures. Drug-regulated ubiquitinylated sites are annotated by their amino acid position within the sequence.
